# Correcting for sparsity and non-independence in glycomic data through a systems biology framework

**DOI:** 10.1101/693507

**Authors:** Bokan Bao, Benjamin P. Kellman, Austin W.T. Chiang, Austin K. York, Mahmoud A. Mohammad, Morey W. Haymond, Lars Bode, Nathan E. Lewis

## Abstract

Glycans are fundamental cellular building blocks, involved in many organismal functions. Advances in glycomics are elucidating the roles of glycans, but it remains challenging to properly analyze large glycomics datasets, since the data are sparse (each sample often has only a few measured glycans) and detected glycans are non-independent (sharing many intermediate biosynthetic steps). We address these challenges with GlyCompare, a glycomic data analysis approach that leverages shared biosynthetic pathway intermediates to correct for sparsity and non-independence in glycomics. Specifically, quantities of measured glycans are propagated to intermediate glycan substructures, which enables direct comparison of different glycoprofiles and increases statistical power. Using GlyCompare, we studied diverse N-glycan profiles from glycoengineered erythropoietin. We obtained biologically meaningful clustering of mutant cell glycoprofiles and identified knockout-specific effects of fucosyltransferase mutants on tetra-antennary structures. We further analyzed human milk oligosaccharide profiles and identified novel impacts that the mother’s secretor-status on fucosylation and sialylation. Our substructure-oriented approach will enable researchers to take full advantage of the growing power and size of glycomics data.

## Introduction

Glycosylation is a highly abundant and complex post-translational modification, decorating between one-fifth and one-half of eukaryotic proteins^1,2^. These diverse carbohydrates account for 12-25% of dry cell mass and have important functional and pathological roles^3,4^. Despite their importance, glycans have complex structures that are difficult to study. The complex structures of glycans arise from a non-template driven synthesis through a biosynthetic network involving dozens of enzymes. A simple change of a single intermediate glycan or glycosyltransferase will have cascading impacts on the final glycans obtained^5,6^. Unfortunately, current data analysis approaches for glycoprofiling and glycomic data lack the necessary systems perspective to easily decode the interdependency of glycans. It is important to understand the network behind the glycoprofiles so that we can better understand the behavior of the process.

New tools aiding in the acquisition and aggregation of glycoprofiles are emerging, making large-scale comparisons of glycoprofiles possible. Advances in mass spectrometry now enable the rapid generation of many glycoprofiles with detailed glycan composition^7–10^, exposing the complex and heterogeneous glycosylation patterns on lipids and proteins^11,12^. Large glycoprofile datasets and supporting databases are also emerging, including GlyTouCan^13^, UnicarbDB^14^, GlyGen and UniCarbKB^15^.

These new technologies and databases provide opportunities to examine global trends in glycan function and their association with disease. However, the rapid and accurate comparison of glycoprofiles can be challenging with the size, sparsity and heterogeneity of such datasets. Indeed, in any one glycoprofile, only a few glycans may be detected among the thousands of possible glycans^16^. Thus, if there is a major perturbation to glycosylation in a dataset, few glycans, if any, may overlap between samples. However, these non-overlapping glycans may only differ in their synthesis by as few as one enzymatic step. Thus, it can be difficult to know which glycans to compare. Furthermore, since glycans often share substantial portions of their biosynthetic pathways with each other, statistical methods that assume independence (e.g., t-tests, ANOVA, etc) are inappropriate for glycomics. Here we address these challenges by proposing glycan substructures, or intermediates, as the appropriate functional units for meaningful glycoprofile comparisons, since each substructure can capture one step in the complex process of glycan synthesis. Thus, using substructures for comparison, we account for the shared dependencies across glycans.

Previous work has investigated the similarity across glycans using glycan motifs, such as, glycan fingerprinting to describe glycan diversity in databases^17^, align glycan structures^18^, identify glycan epitopes in glycoprofiles^19^, deconvolve LC-MS data to clarify glycan abundance^20^, or compare glycans in glycoprofiles leveraging simple structures^21^. These tools use information on glycan composition or epitopes; however, further accounting for shared biosynthetic steps across glycans could provide complete biosynthetic context to all glycan epitopes. That context includes connecting all glycans to the enzymes involved in their synthesis, the order of the enzyme reactions, and information on competition for glycan substrates. Thus, a generalized substructure approach could facilitate the study of large numbers of glycoprofiles by connecting them to the shared mechanisms involved in making each glycan.

Here we present GlyCompare, a method enabling the rapid and scalable analysis and comparison of any number of glycoprofiles, while accounting for the biosynthetic similarities of each glycan. This approach addresses current challenges in sparsity and hidden interdependence across glycomic samples, and will facilitate the discovery of mechanisms underlying the changes among glycoprofiles. We demonstrate the functionality and performance of this approach with both protein-conjugated and unconjugated glycomic analysis, using recombinant erythropoietin (EPO) N-glycosylation and human milk oligosaccharides (HMOs). Specifically, we analyzed sixteen MALDI-TOF glycoprofiles of EPO, where each EPO glycoprofile was produced in a different glycoengineered CHO cell line^9,11^. We also analyzed forty-eight HPLC glycoprofiles of HMO from six mothers^22^. By analyzing these glycoprofiles with GlyCompare, we quantify the abundance of important substructures, cluster the glycoprofiles of mutant cell lines, connect genotypes to unexpected changes in glycoprofiles, and associate a phenotype of interest with substructure abundance and flux. We further demonstrate that such analyses gain statistical power since GlyCompare elucidates and uses shared intermediates. The analysis of the EPO and HMO datasets demonstrate that our novel framework presents a convenient and automated approach to elucidate novel insights into complex patterns in glycobiology.

## Results

### Glycomic data may fail to recover biologically meaningful clusters

Due to the sparsity and non-independence of glycoprofile, clustering and comparing different glycoprofiles can be challenging^23^. We tested this by clustering glycoprofiles from a panel of different Erythropoietin (EPO) glycoforms, each produced in different glycoengineered CHO cell lines. In the clustering, many neighboring samples were not coming from the most genetically similar mutants, and thus did not recapitulate the severity of glycosylation disruption (**Fig. 1a and Supplementary Fig. 1**). These challenges prompted us to develop GlyCompare, a substructure-based approach to glycan analysis. Using GlyCompare, we decomposed the glycoprofiles of glycoengineered EPO into glyco-motif abundance profiles and easily recovered the expected severity of glycoengineered effects (**Fig. 1b**). The glyco-motif abundances mitigate major statistical challenges of working with glycoprofiles. In the next section, we describe how we decompose glycoprofiles into glyco-motif abundance profiles.

**Fig. 1.**
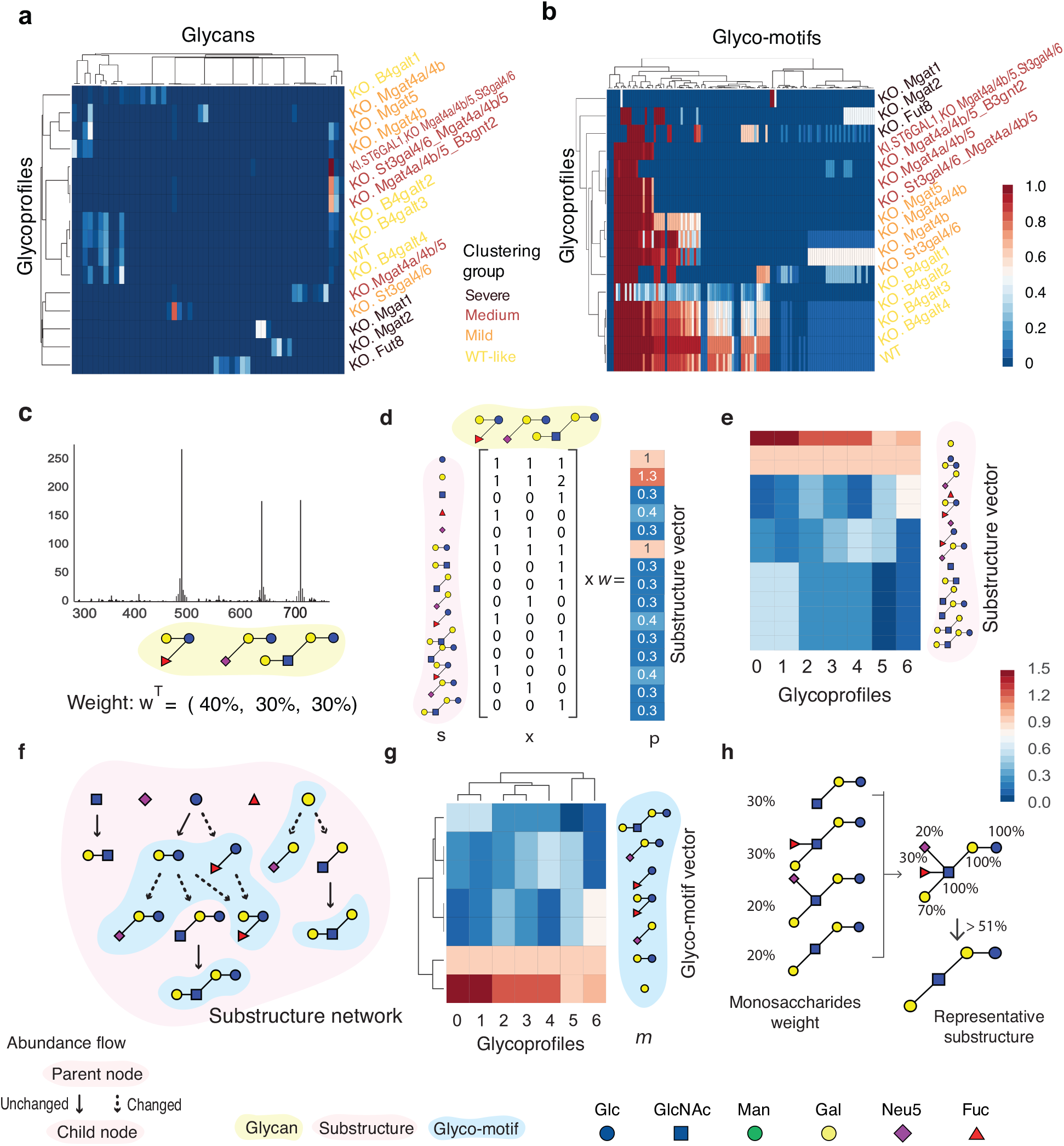
The GlyCompare workflow for glycoprofile decomposition and comparison. **a**, Sixteen glycoprofiles from glycoengineered recombinant EPO cluster poorly when based solely on raw glycan abundance. **b**, GlyCompare was used to compute and cluster EPO glyco-motif vectors, resulting in three dominant clusters of glycoprofiles and a few individuals that have severe changes in their glycan structural pattern (distance threshold=0.5) and twenty-four clusters of glycan substructures (distance threshold=0.19). **c and d**, A glycoprofile with annotated structure and relative abundance is obtained and the glycans are decomposed to a substructure set ***S*** and the presence/absence vectors is built. Presence/absence vectors are weighted by the glycan abundance, and are summed into a substructure vector ***p***. **e**, Seven example glycoprofiles are represented here with their substructure vectors. **f,** To simplify the substructure vectors to contain a minimal number of substructures, a substructure network is constructed to identify the non-redundant glyco-motifs that change in abundance from their precursor substructures. **g**, The glycoprofiles can be re-clustered with simplified glyco-motif vectors for a clearer result. **h**, Clustered substructures can be analyzed to identify the most representative structure in the group. For example, four substructures with different relative abundance were aligned together and the monosaccharides with weight over 51% were preserved.

### GlyCompare decomposes glycoprofiles to facilitate glycoprofile comparison

Glycoprofiles can be decomposed into abundances of glycan intermediate substructures. The resulting substructure profile has richer information than whole glycan profiles and enables more precise comparison across conditions. Since glycan biosynthesis involves long, redundant pathways, the pathways can be collapsed to obtain a subset of substructures while preserving the information of all glycans in the dataset. We call this minimal set of substructures “glyco-motifs.” The GlyCompare workflow consists of several steps wherein glycoprofiles are annotated and decomposed, glyco-motifs are prioritized, and each glyco-motif is quantified for subsequent comparisons. The specific workflow is described as follows.

First, to characterize each glycoprofile with substructures, all substructures in the glycoprofiles are identified and occurrence per glycan is quantified (**Fig. 1c-d**). Thus, a complete set of glycan substructures is obtained for all glycans in all glycoprofiles being analyzed. For each glycoprofile, the abundance of each substructure is calculated by summing the abundance of all glycans containing the substructure. This results in a substructure profile, which stores abundances for all glycan substructures (**Fig. 1e**) in given glycoprofile. The summation over similar structures asserts that similar structures follow the same synthetic paths, which is appropriate for glycosylation wherein synthesis is hierarchical and acyclic (**Supplementary Fig. 2,3**). Therefore, a substructure abundance is not simply a sum over similar structures, it is a meaningful sum over biosynthetic pathways.

**Fig. 2.**
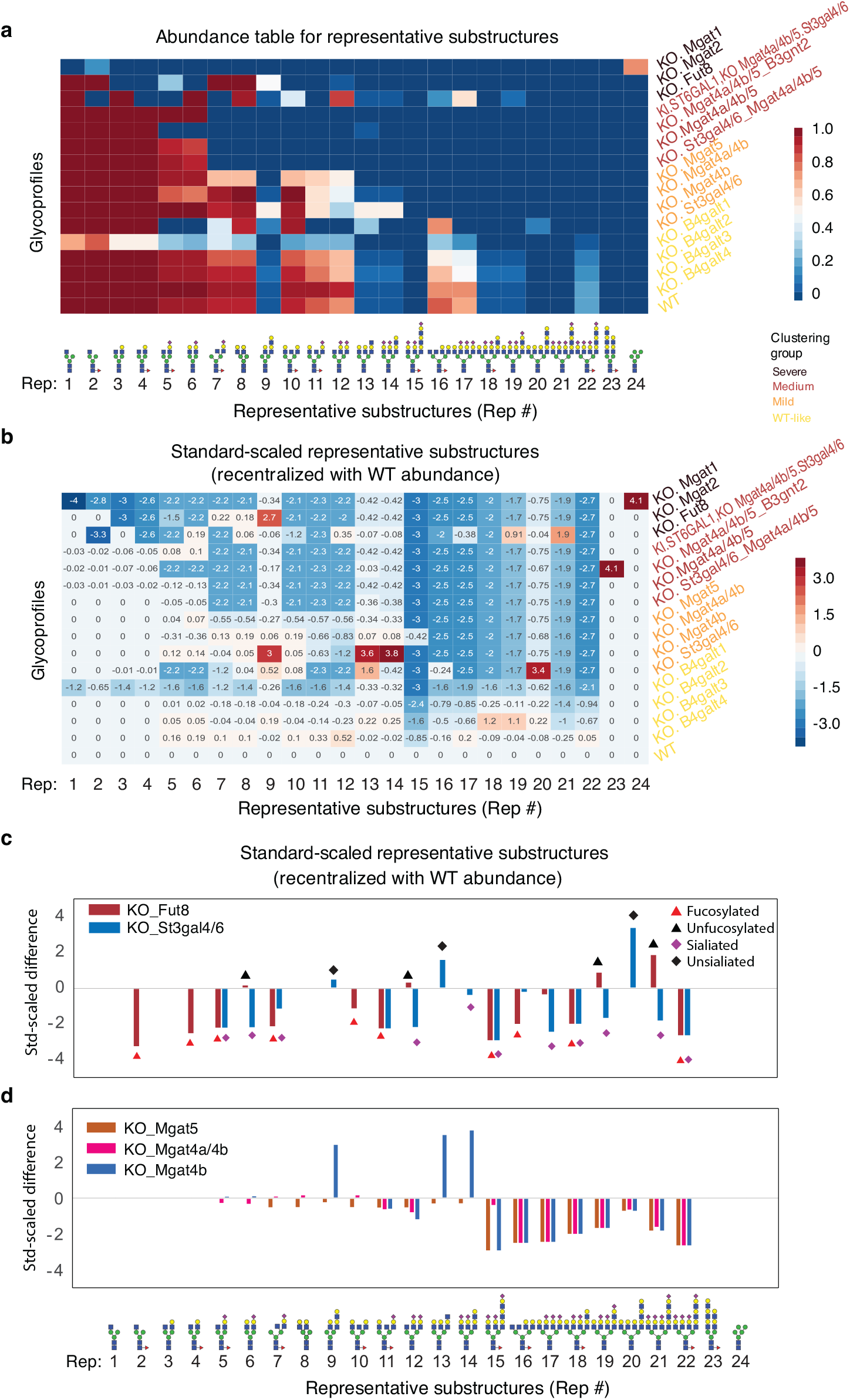
Changes in representative substructures can be quantified and compared to WT. **a**, The representative substructure table contains representative substructures for each of the 24 substructure clusters. The color scale represents the averaged abundances of the substructures in each cluster. The substructures are sorted based on the glycan structure complexity, followed by the number of branches, the degree of galactosylation, sialylation, and fucosylation. **b**, The significantly differentially expressed glycan substructures are illustrated by Standard-scaled abundance of twenty-four glycan substructures, compared with WT. **c**, Differential fucosylation is illustrated for the Fut8 knockout. The red (black) triangles represent the presence/absence of fucose in the representative substructures. Differential sialylation is illustrated for the St3gal4/6 knockout. The purple/black diamonds represent the presence/absence of the sialylation in the representative substructures. **d**, Changes in branching are presented for the Mgat4a/4b/5 knockouts. The tetra-antennary substructures (Rep16 –22) decreased considerably. The triantennary substructures with elongated GlcNac (Rep13 –14) increase significantly (p-value < 0.0031). However, the elongated triantennary structure (Rep15) decreases considerably for the Mgat5 and Mgat4b knockouts, while the Mgat4a/4b knockouts remain high abundance (p-value< 0.0031). In the CHO dataset, the glycan substructure generated by Mgat4a/4b and Mgat5 will be considered as the same topologically.

**Fig. 3.**
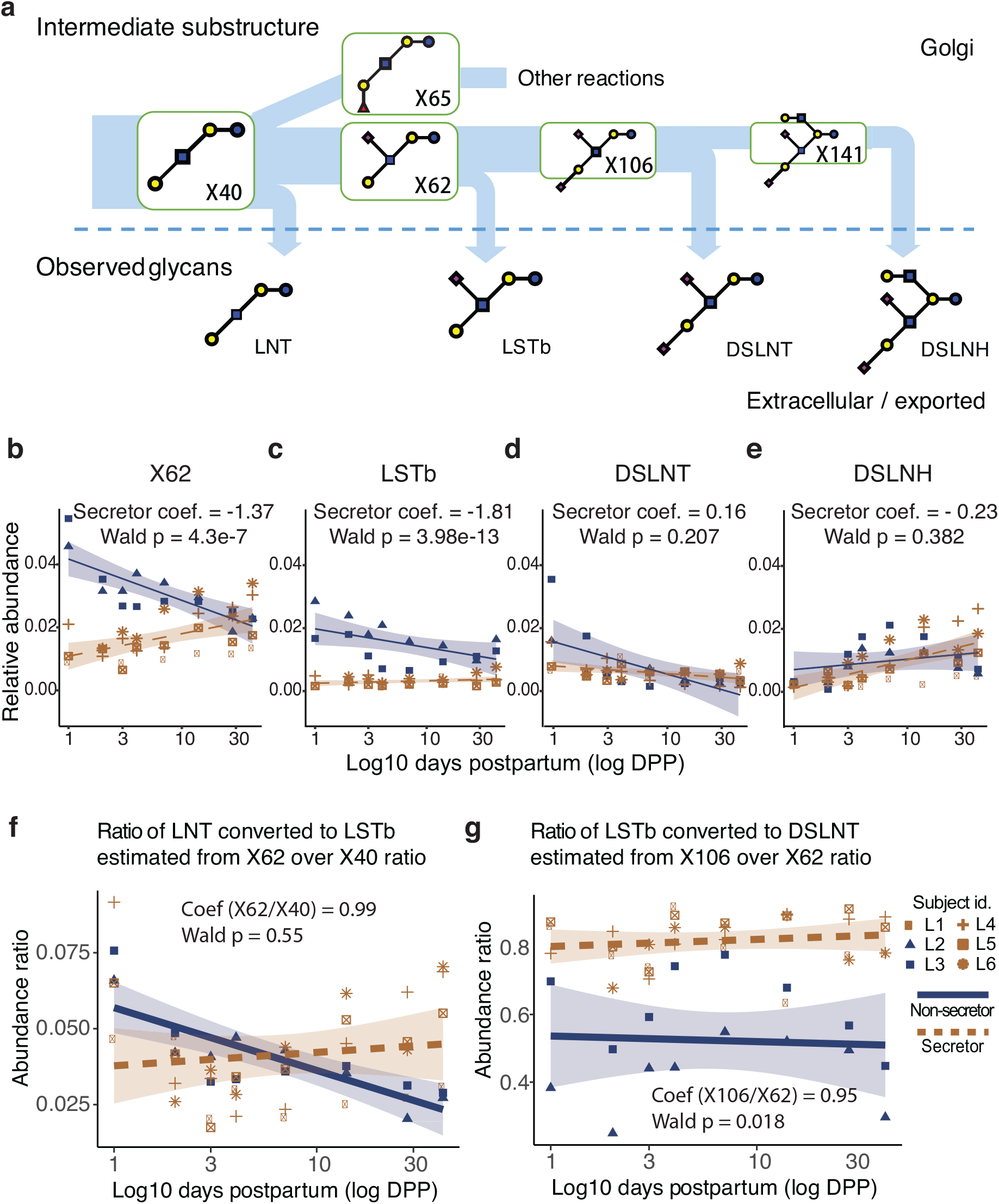
Analysis of intermediate substructures with GlyCompare elucidates associations in abundance and flux with secretor status over time, which are missed in the standard whole-glycan analysis. **a**, The substructure intermediates for four connected HMOs are shown here. The synthesis of larger HMOs must pass through intermediate substructures that are also observed HMOs, where the substructures are as associated with measured HMOs as follow X40=LNT, X62=LSTb, X106=DSLNT, X138=DSLNH. **b-e**, Over time (DPP), X62, LSTb, DSLNT, and DSLNH show different trends for secretors and non-secretors. Furthermore, the abundance of aggregated X62 shows significant positive-correlation with secretor and negative-correlation with non-secretor. **f and g**, Panels examine the product-substrate ratio for two reactions in panel **a**. X40, the LNT substructure, is a precursor to X62, the LSTb substructure, which is a precursor to X106, the DSLNT substructure. We estimate the flux of these conversions from X40 to X62 and X62 to X106 by examining the product-substrate ratio, i.e., the proportion of the total synthesized substrate converted to the product. LSTb/LNT substructure relative abundance ratios are not associated with secretor status while DSLNT/LSTb ratios are. Odds ratios (OR) corresponding the ratio association with secretor status.

Second, to identify the most informative substructures (i.e., glyco-motifs), substructures are prioritized using the substructure network. The substructure network is built by connecting all substructures with biosynthetic steps (**Fig. 1f**). Starting from the monosaccharides, each level of the network represents another biosynthetic step, with one more monosaccharide than the previous level. The edges in the network represent enzymatic additions of each monosaccharide. These edges are weighted by the correlation between the abundances of the substrate and product substructures across all samples. Redundant substructures can be easily identified since their parent-child substructure abundances will be perfectly correlated. Substructure network reduction proceeds by collapsing links with a perfect correlation between substrate and product substructures, and only retaining the product substructure (see methods section for further details). We demonstrate this network reduction in **Fig. 1f**. We identify redundant substructures when the abundance of parent substructures and descendant substructure are perfectly correlated across all glycoprofiles (connected with solid arrow). We remove the parent substructure (substrate) while keeping the child substructure (product). The remaining substructures are termed glyco-motifs; they completely describe the variance at the substructure level. The abundances of all glyco-motifs are then represented as a glyco-motif profile, the minimal subset of meaningful substructure abundances represent glycoprofiles (**Fig. 1f**).

For larger datasets, summarizing the glyco-motifs becomes necessary. Glyco-motif vectors, like glycoprofiles, can be clustered (**Fig. 1g and Supplementary Fig. 4**). We defined a representative substructure as the common structure in a glyco-motif cluster (**Fig. 1h**). The representative substructure describes the glycan features that vary the most across samples. To extract the common structural features in each cluster, we calculated the average weight of each monosaccharide. Monosaccharides with a weight larger than 51% are preserved, which illustrates the predominant structure in the cluster. This allows one to quickly evaluate the distinguishing glycan features that vary across samples in any given dataset.

**Fig. 4:**
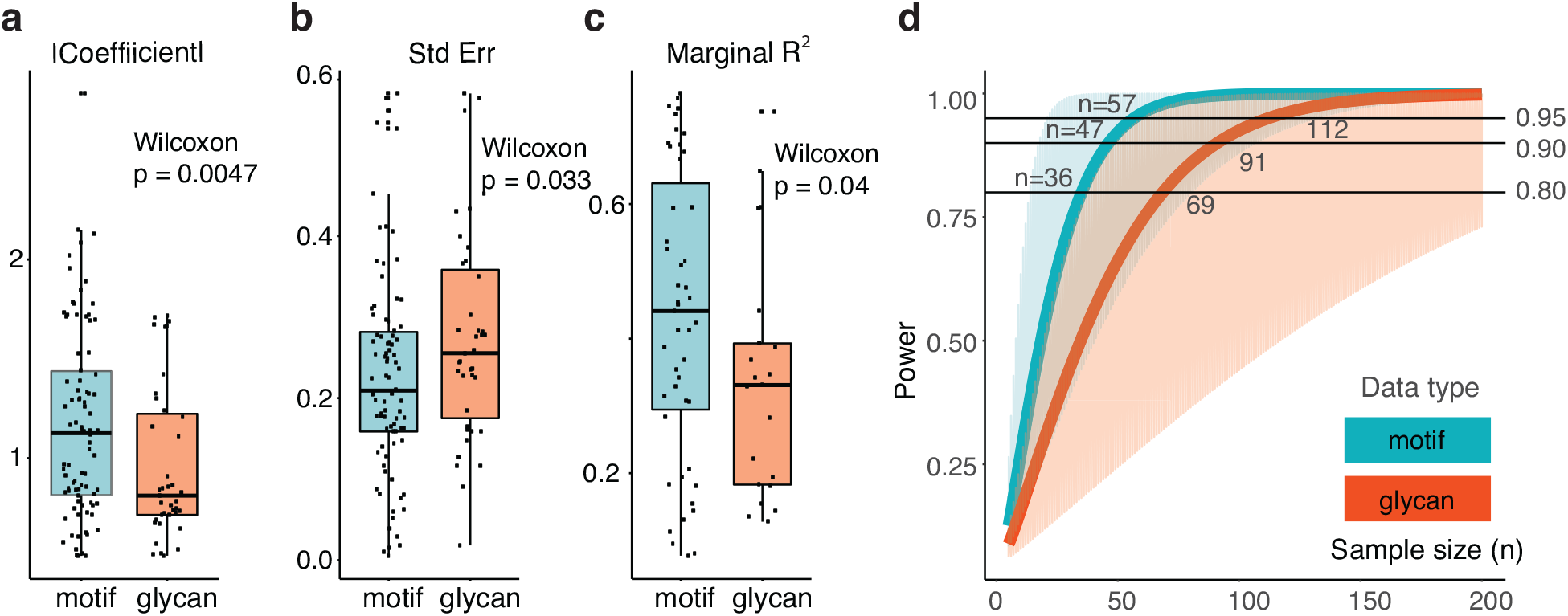
Glyco-motif level statistics require half as many samples to reach the same level of statistical power. **a and b,** The use of glyco-motifs improves measures of regression robustness. The coefficient magnitude and Standard Error indicate the magnitude of the measured effect and the confidence with which a coefficient can be estimated. **c**, The R^2^ describes the effect size of a regression; we used marginal R^2^ (mR^2^) because it was appropriate for the regression models used^34^. **d,** We predicted power for a range of sample sizes (n=5-200) given the median effect size (solid line) within the interquartile range (shaded region) for glyco-motif-trained regressions (mR^2^: median=0.45, Q1=0.31, Q3=0.68) and the median effect size for glycan-trained regressions (mR^2^: median=0.33, Q10.18, Q3=0.44). Here, the use of GlyCompare and glyco-motif abundances required approximately half the number of samples to achieve equivalent power as standard glycan measures.

The workflow we described here successfully connects all glycoprofiles in a data set through their shared intermediate substructures, thus allowing robust analysis of the differences across glycomics samples and the evaluation of the associated genetic bases.

### GlyCompare accurately clusters glycoengineered EPO samples

The poor clustering of the engineered EPO glycosylation data^9^ included clustering of glycoprofiles with low phenotypic similarity (**Fig. 1a and Supplementary Fig. 1,5**). This inconsistency and poor clustering stems from the inherent sparseness of glycoprofiles, i.e., each glycoprofile only has a few glycans. Thus, the matrix of all samples is very sparse, unfit for standard clustering approaches and hard to interpret. Particularly problematic is that pairs of glycans differing in a single monosaccharide are treated as two completely different glycans under standard clustering approaches. Thus, we found that clustering is affected more by the presence or absence of a glycan, rather than structural similarity.

GlyCompare addresses these problems by elucidating hidden similarities between glycans after decomposing glycoprofiles to their composite substructures. The 52 glycans were decomposed into their constituent glycan substructures, resulting in a substructure vector with 613 glycan substructures and a further simplified 120 glyco-motif vector (**Supplementary Fig. 6**). The glyco-motif clustering clearly distinguished the samples based on the structural patterns and separated profiles into groups more consistently associated with the extent of changes in the profile than the raw glycan-based clusters (**Fig. 1b and Supplementary Fig. 5**).

The sixteen glycoprofiles clustered into three groups with a few severely modified outliers (**Fig. 1b**), and the 120 glyco-motifs clustered into twenty-four groups, each summarized by representative substructures Rep1 - Rep24 (**Fig. 2a and Supplementary Fig. 4**). The clusters of glycoprofiles are consistent with the genetic similarities among the host cells. Specifically, the major substructure patterns cluster individual samples into four categories: ‘wild-type (WT)-like’, ‘mild’, ‘medium’ and ‘severe’. The WT-like category contains one group, WT and B4galt1/2/3/4/ knockouts, which contains most of the substructures seen in WT cells. The mild group includes the Mgat4b/4a, Mgat4b, and Mgat5 knockouts, where each lose the tetra-antennary structure, and an St3gal4/6 knockout, which loses the terminal sialylation. The medium category is a group that contains knockouts of St3gal4/6 and Mgat4a/4b/5, knockouts of Mgat4a/4b/5 and B3gnt2, knockouts of Mgat4a/4a/5 with a knock-in of human ST6GAL1, and knockouts of Mgat4a/4b/5 and St3gal4/6. The medium disruption category lost the tri-antennary structure. The ‘severe’ category includes three individual glycoprofiles with knockouts for Fut8, Mgat2, and Mgat1, each of which generate many glycans not detected in the WT-like, mild or medium categories. While some glyco-motif clusters can be seen in the glycoprofile clusters, there are important differences, and the glyco-motif clusters provide more information and improved cluster stability (**Supplementary Fig. 4,7**). These results demonstrate the performance improvement of glyco-motif abundance over glycan abundance in assessing the structural similarity between different glycoprofiles.

### GlyCompare summarizes structural changes across glycoprofiles

GlyCompare helps to more robustly group samples by accounting for the biosynthetic and structural similarities of glycans. Further analysis of the representative structures provides detailed insights into which structural features vary the most across samples. To accomplish this, we rescaled the representative structure abundances and identified significant changes in representative substructure abundances between mutant cells and WT (**Fig. 2a,b**). This highlights the specific structural features of glycans that are impacted when glycoengineering recombinant EPO.

As expected, in the Mgat1 knockout glycoprofile, only high mannose N-glycans are seen. Also, in the Mgat2 knockout, the glycan substructure of bi-antennary on one mannose linkage significantly increases, and the unique structure of bi-antennary LacNac elongated in the N-glycans emerges in the St3gal4/6 and Mgat4a/4b/5 knockouts. Along with expected changes in α-1,6 fucosylation in the Fut8 knockout glycoprofile, we also observed an increase in the tetra-antennary poly-LacNac elongated N-glycan without fucose, which has not been previously reported (One-sided one-sample wilcoxon test, Rep19: p=2.7 × 10^−4^, Rep21: p=2.0 × 10^−4^) (**Fig. 2c**). In the St3gal4/6 knockout (**Fig. 2c**), we observed the relative abundance of structures with sialylation significantly decreased, while the tetra-antennary and triantennary poly-LacNAc elongated N-glycan substructure without sialylation significantly increased (Rep13: p= 1.3 × 10^−3^, Rep20: p=2.3 × 10^−4^). Finally, the Mgat4b, Mgat4a/4b and Mgat5 knockouts (**Fig. 2d**) lose all core tetra-antennary substructures (Rep16: unscaled abundance=0). While triantennary substructures with GlcNac elongation increased significantly for Mgat4b (Rep13: p=2.6 × 10^−3^, Rep14: p=2.5 × 10^−4^), the poly-LacNac elongation structure disappeared. Interestingly, while both the Mgat4b and Mgat5 knockouts do not have the tri-antennary poly-LacNac elongated N-glycan, the Mgat4a/4b mutant keeps a highly abundant poly-LacNac branch (Rep15: p= 2.4 × 10^−4^). Thus, through the use of GlyCompare, we identified the specific glycan features that are impacted not only in individual glycoengineered cell lines, but also features shared by groups of related cell lines.

### GlyCompare reveals phenotype-associated substructures and trends invisible at the whole glycan level

Many secreted and measured glycans are also precursors, or substructures, of larger glycans (**Fig. 3a**). Thus, the secreted and observed abundance of one glycan may not equal to the total amount synthesized. GlyCompare can quantify the total abundance of a glycan by combining the glycan abundance with the abundance of its products. To demonstrate this capability of GlyCompare, we analyzed HMO abundance, and examined the impact of secretor status and days postpartum on HMO abundance. We obtained forty-seven HMO glycoprofiles from 6 mothers (1, 2, 3, 4, 7, 14, 28 and 42 days postpartum (DPP)), 4 “secretor” mothers with functioning FUT2 (α-1,2 fucosyltransferase), and 2 “non-secretor” mothers with non-functional FUT2. With GlyCompare addressing the non-independence of HMOs, we could use powerful statistical methods to study trends in HMO synthesis. Specifically, we used regression models predicting secretor status and DPP from substructure abundance.

We first checked both the glycan-level and substructure-level clustering of the glycoprofile. Samples with same secretor status and days postpartum (DPP) were successfully grouped (**Supplementary Fig. 8**). Further examination of the glyco-motif abundance (i.e., the total amount of substructure synthesized) revealed phenotype-related trends invisible at the level of the whole glycan profile. For example, the LSTb substructure (X62) increased in secretor mothers (Wald p = 2 × 10^−16^) and decreased in non-secretor mothers over time (Wald p < 2 × 10^−16^; **Fig. 3b**). Yet, the same trend was weak or inconsistent for all glycans containing the X62 substructure: LSTb, DSLNT and DSLNH (**Fig. 3b-e**). LSTb weakly shows a similar trend to X62. LSTb decreases over time in non-secretors (Wald p = 6.53 × 10^−4^) but the time-dependent increase in secretors is barely significant (Wald p = 0.046) and the effect size is small (marginal R^2^ = 0.088). Unlike X62, DSLNT shows no significant increase over time (Coef=−0.39, Wald p = 0.17) in secretor mothers. Finally, unlike the decrease over time seen in non-secretors in X62, DSLNH shows a significant increase over time in non-secretors (Wald p = 2.91×10^−8^). The secretor-specific trends in total LSTb are only clearly visible by examining the X62 substructure abundance (**Fig. 3c**). Thus, while secretor status is expected to impact HMO fucosylation, GlyCompare reveals associations with non-fucosylated substructures. Viewing substructure abundance as total substructure synthesized provides a new fundamental measure to the study of glycoprofiles, it also creates an opportunity to explore trends in synthesis.

### GlyCompare identifies flux in HMO biosynthesis

We next applied GlyCompare to explore changes in HMO synthesis over time. For this, we estimate the flux for each biosynthetic reaction by quantifying the abundance ratio of products and substrates from parent-child pairs of glycan substructures. Thus, we could study changes in HMO synthesis through the systematic estimation of reaction flux across various conditions.

We found several reactions strongly associated with secretor status. As expected, the estimated reaction flux from the LNT substructure (X40) to the LNFPI substructure (X65), was strongly associated with secretor status (Wilcox p =1.3 × 10^−12^). In secretors, 36.2% (s.d. 12.7%) of X40 was converted to X65, compared to non-secretors, wherein only 5% (s.d. 1.3%) of X40 was converted.

Although secretor status is defined by the fucosyltransferase-2 genotype, not all secretor-associated reactions were fucosylation reactions. We further explored the secretor-X62 association using the product-substrate ratio to estimate flux. Specifically, we examined the upstream reaction (**Fig. 3f**) of LNT (X40) to LSTb (X62) and the downstream reaction (**Fig. 3g**) of LSTb (X62) to DSLNT (X106). We measured the upstream reaction of LNT converting to LSTb, using the X62/X40 ratio over time, however, no significant change was observed with respect to secretor status (Wald p=0.55). In the conversion of LSTb to DSLNT, we found a secretor-specific reaction increase in flux. Specifically, the X106/X62 ratio was significantly higher (Wald p=0.018) in secretor mothers (**Fig. 4g**; **Supplementary Table 3c**) In the average non-secretor mother, 52.3% (s.d. 15.1%) of LSTb is converted to DSLNT. Meanwhile in secretors, the average conversion rate is 81.8% (s.d. 7.2%). The LSTb to DSLNT conversion rate appears higher in secretors while conversion from the LSTb precursor, LNT, appears unchanged; any changes in sialylation is intriguing, considering secretor status is associated with genetic variation of a fucosyltransferase. Examining the product-substrate ratio has revealed a phenotype-specific reaction propensity thus providing insight to the condition-specific synthesis.

### GlyCompare increases statistical power of glycomics data

GlyCompare successfully provides new insights by accounting for shared biosynthetic routes of measured oligosaccharides. Since it includes information on the similarities between different glycans, we wondered how our approach impacts statistical power in glycan analysis. Thus, to quantify the benefit of glyco-motif analysis, we constructed a large number of regression models associating either glyco-motif abundance or glycan abundance, with a DPP and secretor status (see **Methods**). We found that regressions trained with glyco-motif abundance are more robust than those trained on whole glycan HMO abundance, as indicated by the increased coefficient magnitude (Wilcoxon p = 0.0047, **Fig. 4a**), and decreased standard error (Wilcoxon p = 0.033, **Fig. 4b**). An increase in the stability of a statistic can result in an increased effect size. Consistent with the increased coefficient magnitude and decreased standard error, the effect size also increased, as measured by the marginal R^2^ (mR^2^) of glyco-motif-trained regressions (Wilcoxon p=0.04, **Fig. 4c**). These effects were confirmed with a bootstrapping t-test; bootstrapping p-values were less than or equal to Wilcoxon p-values within 0.001. Increases in statistic magnitude, statistic stability, and effect size are all expected to increase the power of an analysis. Using the median, 1^st^ quartile, and 3^rd^ quartile of observed mR^2^, we estimated the expected power of glyco-motif-trained and glycan-trained regressions at various sample sizes. The expected power of a glyco-motif-trained regression reaches 0.8 at 36 samples and 0.9 with 57 samples while a glycan-trained regression requires more than double the sample size to reach a comparable power (**Fig. 4d**). Thus, using GlyCompare for glyco-motif-level analysis can substantially increase the robustness and statistical power in glycomics data analysis since it allows for the comparison of different glycans who share biosynthetic steps.

## Discussion

Glycosylation has generally been studied from the whole-glycan perspective using mass spectrometry and other analytical methods. From this perspective, two glycans that differ by only one monosaccharide are distinct and are not directly comparable. Thus, the comparative study of glycoprofiles has been limited to changes between glycans shared by multiple glycoprofiles or small manually curated glycan substructures^17^. GlyCompare sheds light on the hidden biosynthetic interdependencies between glycans by integrating the biosynthetic pathways into the comparison. Glycoprofiles are converted to glyco-motif profiles, wherein each substructure abundance represents the cumulative abundance of all glycans containing that substructure. This enumeration and quantification of substructures can be easily scaled up to include many glycoprofiles in large datasets. Additionally, since no prior information is required beyond glycan identities and quantities, the method can even facilitate analysis of glycans with limited characterization. Thus, it brings several advantages and new perspectives to enable the systematic study of glycomics data.

First, the GlyCompare platform computes a glyco-motif profile (i.e., the abundances of the minimal set of glycan substructures) that maintains the information of the original glycoprofiles, while exposing the shared intermediates of measured glycans. These sample-specific glyco-motif profiles more accurately quantify similarities across glycoprofiles. This is made possible since glycans that share substructures also share many biosynthetic steps. If the glycan biosynthetic network is perturbed, all glycans synthesized will be impacted and the nearest substructures will directly highlight where the change occurred. For example, in EPO glycoprofiles studied here, the tetra-antennary structure is depleted in the Mgat4a/4b/5 knockout group and the downstream sialylated substructure depleted when St3gal4/6 were knocked out. Such structural patterns emerge in GlyCompare since the tool leverages shared intermediate substructures for clustering, thus identifying common features in glycans measured across diverse samples.

Second, new trends in glycan biosynthetic flux become visible at the substructure level. For example, in the HMO data set, multiple HMOs are made through a series of steps from LNT to DSLNH (**Fig. 4a**). Only when the substructure abundances and product-substrate ratios are computed are we able to observe the secretor-dependent differences in the abundance of the LSTb substructure, X62. This is particularly interesting since secretor status is defined by changes in α-1,2 fucosylation, but we see here additional secretor-dependent changes to sialylated structures with no fucose. These are the systemic effects invisible without a systems-level perspective due to the interconnected nature of glycan synthesis; this disparity underlines the power of this method.

Third, the sparse nature of glycomic datasets and the synthetic connections between glycans make glycomic data unfit for many common statistical analyses. However, the translation of glycoprofiles into substructure abundance provides a framework for more statistically powerful and robust analysis of glycomic datasets. Single sample perturbations, such as the knockouts in the glycoengineered EPO, can be compared to wild-type; all substructure data can be normalized and then rigorously distinguished from the control using a one sample Wilcoxon-test. Furthermore, conditions or phenotypes with many glycoprofiles, such as the secretor status in the HMO dataset, can be compared using a variety of statistical methods to evaluate the association between the phenotypes and glycosylation. For example, in HMO data, we revealed that the α-1,2 fucose substructure is enriched in secretor status, consistent with the previous studies^24–26^. Because the substructure approach includes comparisons of glycans that are not shared across the different samples, but that share intermediates, GlyCompare decreased sparsity and increased statistical power. Thus, one can obtain richer glycan comparisons of representative substructures, total synthesized abundance, and flux.

Finally, in combination with the substructure network, we can systematically study glycan synthesis. The product-substrate ratio provides an estimation of flux through the glycan biosynthetic pathways. Using the HMO dataset, we demonstrate the power of this perspective by showing that more LSTb is converted to DSLNT in the secretor mother. The perspectives made available through GlyCompare are not limited to Wilcoxon-tests and regression models. Because the substructure-level perspective minimizes biosynthetic dependency between glycans, glyco-motif abundances can be used with nearly any statistical model or comparison demanded by a dataset. We have reduced the sparse and non-independent nature of glycoprofiles, thereby making countless comparisons and new analyses possible.

## Conclusions

In conclusion, GlyCompare provides a novel paradigm for describing complex glycoprofiles, thus enabling a wide range of analyses and facilitating the acquisition of detailed insights into the molecular mechanisms controlling all types of glycosylation.

## Acknowledgements

Many thanks to Joshua Klein for providing guidance on using glypy (https://github.com/mobiusklein/glypy). Thanks to Philip Gordts for insightful comments. This work was conducted with support from the Novo Nordisk Foundation provided to the Center for Biosustainability at the Technical University of Denmark (NNF10CC1016517: A.W.T.C.), NIGMS (R35 GM119850: B.B., B.K.P., N.E.L.), NICHD (R21 HD080682: L.B.) and USDA (USDA/ARS 6250-6001; M.W.H). This work is a publication of the U.S. Department of Agriculture/Agricultural Research Service, Children’s Nutrition Research Center, Department of Pediatrics, Baylor College of Medicine, Houston, Texas. The contents of this publication do not necessarily reflect the views or policies of the U.S. Department of Agriculture, nor does mention of trade names, commercial products, or organizations imply endorsement from the U.S. government.

## Author contributions

B.B, B.P.K. designed the work. B.B., B.P.K., A.W.T.C., A.K.Y., and N.E.L. performed data analysis. M.A.M., M.W.H., and L.B. provided HMO data. The manuscript was written by B.B., B.P.K., A.W.T.C., L.B., and N.E.L.

## Methods

### Data, source code, examples, Jupyter notebooks for generating manuscript figures, and CodeOcean capsule available at

https://github.com/LewisLabUCSD/GlyCompare

### N-glycosylation of EPO glycoprofile collection and analysis

N-glycosylation data were previously published and described elsewhere^9^. Briefly, these data were generated as follows. Different combinations of glycosyltransferase genes were knocked out using zinc-finger nucleases. Both single gene and multigene mutants were generated. Erythropoietin (EPO) was transfected into the library of glycoengineered cell lines. After overexpression of EPO, glycans were cleaved using PNGase, and then assayed by mass spectrometry. Upon retrieval of these data from the study, we picked 16 glycoprofiles that are used again in their following up study ^11^ and further processed the data as follows. All measurements were taken from distinct samples.

Glycan substructures were extracted from the observed glycans. Substructure abundance was calculated from glycan abundance of all glycans containing the substructure. A minimal set of 120 glyco-motifs substructures identified by substructure network to compare the mutants. Finally, representative substructures were extracted to pool abundance and summarize the structural changing across mutants. Each of these operations is further specified below.

### HMO glycoprofile collection and analysis

Following Institutional Review Board approval (Baylor College of Medicine, Houston, TX), lactating women were given written informed consent. Women with diabetes or impaired glucose tolerance, anemia, or renal or hepatic dysfunction were excluded from the study. Women were 18-35 years of age, had uncomplicated singleton pregnancies with vaginal delivery at term (>37 weeks) and pregnancy Body Mass Index (BMI) remained <26kg/m2. Infants were healthy and exclusively breastfed. Forty-eight milk samples were collected from 6 human mothers (1, 2, 3, 4, 7, 14, 28, and 42 days postpartum (DPP)). More information on subject selection, exclusion, study design, and breast milk collection has already been published ^22,27^

HMO composition and abundance was measured by high-performance liquid chromatography (HLPC) following fluorescent derivatization with 2-aminobenzamide (2AB, CID: 6942) as previously described ^28,29^. Raffinose (CHEBI:16634, CID:439242), a non-HMO oligosaccharide, was added to each milk sample as an internal standard at the very beginning of sample preparation to allow for absolute quantification. Of the 300-500 predicted HMO, the 16 most abundant HMO were detected based on retention time comparison with commercial standard oligosaccharides and mass spectrometry analysis including 2-fucosyllactose (2’FL), 3-fucosyllactose (3’FL), 3-sialyllactose (3’SL), lacto-N-tetrose (LNT), lacto-N-neotetraose (LNnT), lacto-N-fucopentaose (LNFP1, LNFP2 and LNFP3), sialyl-LNT (LSTb and LSTc), difucosyl-LNT (DFLNT), disialyllacto-N-tetraose (DSLNT), fucosyl-lacto-N-hexaose (FLNH), difucosyl-lacto-N-hexaose (DFLNH), fucosyl-disialyl-lacto-N-hexaose (FDSLNH) and disialyl-lacto-N-hexaose (DSLNH). Because these are the most abundant HMOs, these glycoprofiles represent the least sparse subset of the entire HMO glycoprofile which is extremely sparse. GlyTouCan IDs for each HMO are listed in **Supplementary Table 2**. Technicians were blinded to metadata associated with each sample. In addition to absolute concentrations, the proportion of each HMO per total HMO concentration (sum of all integrated HMO) was calculated and expressed as relative abundance (% of total, *w*_*i*_/*Σw*_∗_). The presence of 2-FL defines secretor status. All measurements were taken from distinct samples.

HMO abundances profiles were treated similarly to the N-glycans. We identified and quantified 26 glyco-motifs from 121 substructures. We compared glyco-motif abundance and their abundance ratios directly to secretor status along the log of days postpartum.

### Glycoprofile preprocess procedures

Three procedures were used for preprocessing the studied glycoprofiles (**Fig. 1c**). First, glycoprofiles are parsed into glycans with abundance. In each glycoprofile, the glycans are manually drawn and exported with GlycoCT format using the GlyTouCan Graphic Input tool^13^. GlycoCT formatted glycans are loaded into Python (version 3+) and initialized as glypy.glycan objects using the *glypy* (version 0.12.1). Assuming we have a glycoprofile ***i***, the corresponding abundance of each glycan ***j*** in glycoprofile ***i*** is represented by *g*_*ij*_. For example, the relative m/z peak in the mass spectrum or the abundance value in an HPLC trace, is calculated relative to the total abundance of glycans in this glycoprofile *g*_*ij*_/*Σg*_*i*∗_. Glycans with ambiguous topologies are handled by assuming they belong to every possible structure with equal probability, thereby creating all possible ***n*** structures but with *g*_*ij*_/*nΣg*_*i*∗_ abundance of each. Second, glycans are annotated with glycan substructure information, and this information is transformed into the substructure vector. Substructures within a glycan are exhaustively extracted by breaking down each linkage or a combination of linkages of the studied glycan. Note that this method cannot currently deal with glycans with ring topology. All substructures extracted are merged into a substructure set ***S***. Substructures are sorted by the number of monosaccharides and duplicates are removed. Then, each glycan is matched to the substructure set ***S*** producing a binary glycan substructure presence (1) or absence (0) vector, *x*_*ij*_. Lastly, a substructure (abundance) vector is calculated as *p*_i_ = *Σx*_*ij*_*g*_*ij*_/*Σg*_*i*∗_ representing the abundance of the substructures ***s*** in this glycoprofile, where *p*_*i*_ = (*s*_1*i*_,…,*s*_*ni*_). Third, a substructure network is built based on the substructure vectors. The substructure network is a directed acyclic graph wherein each node denotes a glycan substructure. Given the substructure set S, the root node starts from the monosaccharides or a defined root core structure, and a child node is a substructure that has only one monosaccharide added to its parent node. We note that one child node might have multiple parent nodes and vice versa. The child node depends on its parent node(s) since it cannot exist alone without any parent node.

### Generating the glyco-motif vector bases on the substructure abundance

A larger subset of the substructure network is necessary to uniquely describe a more diverse set of glycoprofiles while fewer substructures are needed to describe more similar glycoprofiles sufficiently. Comparisons become more focused when only examining these variable substructures. By checking the substructure network, the substructures that have the same abundance can be merged without any information loss. In other words, after the substructure network is generated, it is simplified by merging the substructure nodes. As illustrated in **Fig. 1f**, the parent-child substructure pairs with perfectly correlated abundance (solid arrow), can be merged. We remove the parent node while keeping the child node. Furthermore, an epitope substructure can also be removed if they are 100% correlated with the bigger substructure containing that epitope. Base on our rule, the merging criteria are based on how child substructure node *S*_*b*_ depends on the parent substructure node *S*_*a*_. The dependency is the Pearson correlation of their abundance across all glycoprofiles, *corr*(*s*_*a*∗_,*S*_*b*∗_). If the correlation is 1, we can conclude that the addition of the specific monosaccharide is not perturbed across all glycoprofiles, which means they carry the same information. Thus, the parent node can be pruned without information loss. All remaining nodes, namely, the glyco-motifs, are used to cluster the glycoprofiles.

Meanwhile, we use the “monosaccharides weight” to track the nodes merging process. All node weights are initialized as 1. When a node is removed, the weight is equally divided and distributed to child nodes whose correlation with the removed node is 1. Since this method redistributes weight from the root to leaves, the last decedent substructure node with a non-unique abundance pattern gains the most weight. The weights **W** are used later for generating the representative substructures.

### Procedures for glycoprofile clustering and identifying representative glycan substructures

The preprocessed glycoprofiles (see details in the “glycoprofile preprocess procedures”) generate the substructure vectors to enable further clustering analysis. Here we used the Pearson correlation and ‘complete’ distance to cluster the glycoprofiles. This procedure clusters the glycoprofiles and substructures.

To identify the representative glycan substructures, a set of glycan substructures with weights ***W*** are first aligned. Then, we calculate the sum of monosaccharide weights for each glycan substructure. The representative substructure is thus defined as the glycan substructures with their summed monosaccharide weights greater than 51% of the total weight of glycan substructures. Lastly, the averaged abundances of the representative substructures are generated to assess their differential expressions between different glycoprofiles.

### Test the abundance changes on representative substructures

We use the representative substructures to summarize and analyze the structural and quantitative changes across glycoprofiles. For the abundance of a representative substructure in a glyco-motif cluster, we use the substructure monosaccharide weights to calculate the weighted average of substructure abundance. Since the abundance range of representative substructures across different glycoprofiles are different, we re-centralized the representative substructure abundance based on WT and scaled them with standard deviation. We can find many interesting signals since there are many representative substructures extremely deviating from the WT’s abundance. Since the abundance distributions are not normally distributed, we used a one-sided 1-sample Wilcoxon test to test if the abundance of a representative substructure in a glycoprofile is significantly divergent. Effect size, r, was calculated as z/sqrt(N)^30^. A Bonferroni correction (n=16) was used to correct for multiple testing, so p=0.0031 is used as criteria and effect sizes are all above 0.68.

### Testing the substructure-phenotype association

We estimated the influence of Secretor status on HMO and glyco-motif abundance using generalized estimating equation (GEE, R3.6∷geepack^31,32^). GEE models account for resampling bias in longitudinal measurements^33^; other regression models, like generalized linear models, overestimate the sample size and power by ignoring this bias. Unlike mixed effect models, which can account for resampling bias, GEE allows non-linear relations between the outcome and covariates, while accounting for correlation among repeated measurements from the same subject. Here we used GEE with exchangeable correlation structure (assuming the within-subject correlation between any two time-points is ρ). To stabilize the variance and equalize the range, we log and z-score standardized each HMO and glyco-motif measurement. We also used the log of days postpartum (DPP) to linearize the relationship over time. The Wald test was used to measure the significance of Secretor status contribution. For additional information and diagnostic statistics for specific regressions, see **Supplementary Table 3a,b**. All regression can be found in **Supplementary Fig. 9.**

### Product-substrate ratio as a proxy for flux and estimating flux-phenotype associations

To further isolate glyco-motif-specific effects from biosynthetic biases, we explored methods to control for the product-substrate relations. First, we isolated the relative abundance of parent-child pairs of glyco-motifs in the substructure network; these are product-substrate relations like LNT and LSTb. Glyco-motif abundance represents the total substructure synthesized; therefore, when we examine the product-substrate ratio, we measure the total amount of the substrate substructure converted to the product substructure in the sample. Thus, the product-substrate ratio is a proxy for flux. Using logistic GEE regression modeling, similar to the approach used for testing substructure-phenotype associations, we can measure the influence of estimated flux between two glycans on secretor status; here we predicted secretor status from estimated flux log(DPP). For additional information and diagnostic statistics, see **Supplementary Table 3c.**

### Glyco-motif Abundance Robustness and Power Analysis

GEE models, similar to those used in **Supplementary Fig. 9**, were trained using either glyco-motif or whole HMO relative abundance. To stabilize the variance, equalize the range and make the regressions comparable, we used a square root and z-score normalization on each HMO and glyco-motif measurement. Glyco-motif or glycan relative abundance was predicted from either DPP alone, Secretor status alone, DPP + Secretor status, or DPP + Secretor status + DPP:Secretor. To avoid biasing the analysis with misfit or uninformative models, models with small coefficients (|coef|<0.5) or extremely non-normal abundance distributions (Shapiro-Wilks p < 0.001) were removed. Model robustness measures including, coefficient magnitude (n_glycan-stats_=39, n_motif-stats_=86), standard error (n_glycan-stats_=39, n_motif-stats_=86) and marginal R^2^ (n_glycan-stats_=21, n_motif-stats_=47) were used to compare model performance. Robustness measures from glycan-trained and glyco-motif-trained models were compared using one-sided Wilcoxon rank sum test with continuity correction. We validated these findings using a 10,000 iteration one-sided, two-sample bootstrapping t-tests (Rv3.6∷nonpar∷boot.t.test); bootstrapping p-values were less than or equal to Wilcoxon rank sum p-values within 0.001. Finally, using the Rv3.6∷pwr∷pwr.r.test v1.2.2 package, statistical power was predicted between n=5 and n=200 for the median and interquartile range of effect sizes observed in glyco-motif-trained and glycan-trained models.

## References

1. Khoury, G. A., Baliban, R. C. & Floudas, C. A. Proteome-wide post-translational modification statistics: frequency analysis and curation of the swiss-prot database. Sci. Rep. 1, (2011).

2. Apweiler, R., Hermjakob, H. & Sharon, N. On the frequency of protein glycosylation, as deduced from analysis of the SWISS-PROT database. Biochim. Biophys. Acta 1473, 4–8 (1999).

3. RodrÍguez, E., Schetters, S. T. T. & van Kooyk, Y. The tumour glyco-code as a novel immune checkpoint for immunotherapy. Nat. Rev. Immunol. 18, 204–211 (2018).

4. Gutierrez, J. M. et al. Genome-scale reconstructions of the mammalian secretory pathway predict metabolic costs and limitations of protein secretion. bioRxiv 351387 (2018). doi:10.1101/351387

5. Gabius, H.-J., André, S., Kaltner, H. & Siebert, H.-C. The sugar code: functional lectinomics. Biochimica et Biophysica Acta (BBA)-General Subjects 1572, 165–177 (2002).

6. Spahn, P. N. & Lewis, N. E. Systems glycobiology for glycoengineering. Curr. Opin. Biotechnol. 30, 218–224 (2014).

7. Holst, S. et al. N-glycosylation Profiling of Colorectal Cancer Cell Lines Reveals Association of Fucosylation with Differentiation and Caudal Type Homebox 1 (CDX1)/Villin mRNA Expression. Mol. Cell. Proteomics 15, 124–140 (2016).

8. Reiding, K. R., Blank, D., Kuijper, D. M., Deelder, A. M. & Wuhrer, M. High-throughput profiling of protein N-glycosylation by MALDI-TOF-MS employing linkage-specific sialic acid esterification. Anal. Chem. 86, 5784–5793 (2014).

9. Yang, Z. et al. Engineered CHO cells for production of diverse, homogeneous glycoproteins. Nat. Biotechnol. 33, 842–844 (2015).

10. Anugraham, M. et al. Specific glycosylation of membrane proteins in epithelial ovarian cancer cell lines: glycan structures reflect gene expression and DNA methylation status. Mol. Cell. Proteomics 13, 2213–2232 (2014).

11. Čaval, T., Tian, W., Yang, Z., Clausen, H. & Heck, A. J. R. Direct quality control of glycoengineered erythropoietin variants. Nat. Commun. 9, 3342 (2018).

12. Riley, N. M., Hebert, A. S., Westphall, M. S. & Coon, J. J. Capturing site-specific heterogeneity with large-scale N-glycoproteome analysis. Nat. Commun. 10, 1311(2019).

13. Aoki-Kinoshita, K. et al. GlyTouCan 1.0--The international glycan structure repository. Nucleic Acids Res. 44, D1237–42 (2016).

14. Campbell, M. P. et al. Validation of the curation pipeline of UniCarb-DB: building a global glycan reference MS/MS repository. Biochim. Biophys. Acta 1844, 108–116 (2014).

15. Campbell, M. P. et al. UniCarbKB: building a knowledge platform for glycoproteomics. Nucleic Acids Res. 42, D215–21 (2014).

16. Cummings, R. D. The repertoire of glycan determinants in the human glycome. Mol. Biosyst. 5, 1087–1104 (2009).

17. Rademacher, C. & Paulson, J. C. Glycan fingerprints: calculating diversity in glycan libraries. ACS Chem. Biol. 7, 829–834 (2012).

18. Hosoda, M. et al. MCAW-DB: A glycan profile database capturing the ambiguity of glycan recognition patterns. Carbohydr. Res. 464, 44–56 (2018).

19. Alocci, D. et al. Understanding the glycome: an interactive view of glycosylation from glycocompositions to glycoepitopes. Glycobiology 28, 349–362 (2018).

20. Klein, J., Carvalho, L. & Zaia, J. Application of network smoothing to glycan LC-MS profiling. Bioinformatics 34, 3511–3518 (2018).

21. Sharapov, S. et al. Defining the genetic control of human blood plasma N-glycome using genome-wide association study. bioRxiv 365486 (2018). doi:10.1101/365486

22. Mohammad, M. A., Hadsell, D. L. & Haymond, M. W. Gene regulation of UDP-galactose synthesis and transport: potential rate-limiting processes in initiation of milk production in humans. Am. J. Physiol. Endocrinol. Metab. 303, E365–76 (2012).

23. Ashwood, C., Pratt, B., MacLean, B. X., Gundry, R. L. & Packer, N. H. Standardization of PGC-LC-MS-based glycomics for sample specific glycotyping. Analyst 144, 3601–3612 (2019).

24. Koda, Y., Soejima, M., Liu, Y. & Kimura, H. Molecular basis for secretor type alpha(1,2)-fucosyltransferase gene deficiency in a Japanese population: a fusion gene generated by unequal crossover responsible for the enzyme deficiency. Am. J. Hum. Genet. 59, 343–350 (1996).

25. Kudo, T. et al. Molecular genetic analysis of the human Lewis histo-blood group system. II. Secretor gene inactivation by a novel single missense mutation A385T in Japanese nonsecretor individuals. J. Biol. Chem. 271, 9830–9837 (1996).

26. Viverge, D., Grimmonprez, L., Cassanas, G., Bardet, L. & Solere, M. Discriminant carbohydrate components of human milk according to donor secretor types. J. Pediatr. Gastroenterol. Nutr. 11, 365–370 (1990).

27. Mohammad, M. A. & Haymond, M. W. Regulation of lipid synthesis genes and milk fat production in human mammary epithelial cells during secretory activation. Am. J. Physiol. Endocrinol. Metab. 305, E700–16 (2013).

28. Bode, L. et al. Human milk oligosaccharide concentration and risk of postnatal transmission of HIV through breastfeeding. Am. J. Clin. Nutr. 96, 831–839 (2012).

29. Alderete, T. L. et al. Associations between human milk oligosaccharides and infant body composition in the first 6 mo of life. Am. J. Clin. Nutr. 102, 1381–1388 (2015).

30. Rosenthal, R. & Rubin, D. B. Further issues in effect size estimation for one-sample multiple-choice-type data. Psychological Bulletin 109, 351–352 (1991).

31. Yan, J. & Fine, J. Estimating equations for association structures. Stat. Med. 23, 859–74; discussion 875–7,879–80 (2004).

32. Halekoh, U., Højsgaard, S., Yan, J. & Others. The R package geepack for generalized estimating equations. J. Stat. Softw. 15, 1–11 (2006).

33. Zeger, S. L. & Liang, K. Y. Longitudinal data analysis for discrete and continuous outcomes. Biometrics 42, 121–130 (1986).

34. Zheng, B. Summarizing the goodness of fit of generalized linear models for longitudinal data. Stat. Med. 19, 1265–1275 (2000).

