## Supplementary for "Correcting for sparsity and non-independence in glycomic data through a systems biology framework"

**Supplementary Table 1 – Glossary of analyses terms**

**Substructures types examined in each section**

| Results section | Substructure | Explanation |
|---|---|---|
| GlyCompare decomposes glycoprofiles to facilitate glycoprofile comparison | Glyco-motif | EPO clustering was done with glyco-motif abundance |
| GlyCompare decomposes glycoprofiles to facilitate glycoprofile comparison | All | Overview of methods discusses every substructure type. |
| GlyCompare accurately clusters glycoengineered EPO samples | Glyco-motif | EPO clustering was done with glyco-motif abundance |
| GlyCompare summarizes structural change across glycoprofiles | Representative substructure | EPO clusters were examined for enrichment and depletion of representative substructures |
| GlyCompare reveals phenotype-associated substructures and trends invisible at the whole glycan level | Substructure | All HMO substructures were used to avoid merging substructure matching known HMOs. This was necessary to allow comparison to know structures |
| GlyCompare identifies condition-specific synthesis dynamics | Substructure | All HMO substructures were used to avoid merging substructure matching known HMOs. This was necessary to allow comparison to know structures |
| GlyCompare increases statistical power of glycomics data | Glyco-motif | Just HMO glyco-motifs were used to avoid artificially overpowering the analysis. |

**Supplementary Table 2 | HMO abbreviations**

HMO abbreviations are specified in this table. Complete GlycoCT structures and GlyTouCan accession, for all HMO and EPO glycans used in this study, can be accessed at https://github.com/LewisLabUCSD/GlyCompare/blob/master/example_data/Glycan_Structures.complete.xlsx

| HMO | Abbreviation | GlyTouCan Accession |
|---|---|---|
| LNT | Lacto-N-tetrose | G45827GY |
| LNnT | Lacto-N-neotetrose | G48059CD |
| 2’FL | 2’-fucosylactose | G10422IZ |
| 3FL | 3-fucosylactose | G06210XB |
| 3’SL | 3’-sialyllactose | G91237TK |
| LNFPI | Lacto-N-fucopentose I | G01650PH |
| LNFPII | Lacto-N-fucopentose II | G98173LG |
| LNFPIII | Lacto-N-fucopentose III | G83916HL |
| LSTb | LS-tetrasaccharide b | G19017MP |
| LSTc | LS-tetrasaccharide c | G72506RN |
| DSLNT | Disialyllactose-N-tetrose | G38710SX |
| DFLNT | Difucosyllacto-N-tetrose | G70115XG |
| FLNH | Fucosyllacto-N-hexose | G24504JY |
| DSLNH | Disialyllacto-N-hexaose | G47928KI |
| DFLNH | Difucosyllacto-N-hexaose | G63053GR |
| FDSLNH | Fucodisialyllacto-N-hexaose | N/A |

**Supplementary Table 3 | Complete information on generalized estimating equation models**

The tables below specify the coefficient summary, confidence intervals, Wald test p-values. We report general model statistics including number of observations and groups, and degrees of freedom. We report effect size with marginal correlation for gaussian regressions and entropy for logistic regression (Zhang, 2000). Finally, for gaussian regressions we report the Shapiro-Wilk's p-value for normality of a distribution.

**a.** Gaussian GEE, predicting motif abundance from secretor status while controlling for DPP

$GEE(z(log(\mathbf{X62}+\epsilon)) \sim log(DPP) + Secretor$,id=subject,corstr='exchangeable')

| | Coef | 95% CI | Pr(Wald) | N. Obs | N. Groups | Marginal $R^2$ | df | Pr(Shapiro-Wilks) |
|---|---|---|---|---|---|---|---|---|
| (Intercept) | 0.74030 | (0.3485 - 1.132) | 6.118e-03 | 47 | 6 | 0.45 | 44 | 0.53 |
| Secretor | -1.36800 | (-0.6422 - -2.095) | 4.329e-07 | | | | | |
| log(DPP) | 0.09137 | (0.06535 - 0.1174) | 0.5294 | | | | | |

$GEE(z(log(\mathbf{LSTb}+\epsilon)) \sim log(DPP) + Secretor$,id=subject,corstr='exchangeable')

| | Coef | 95% CI | Pr(Wald) | N. Obs | N. Groups | Marginal $R^2$ | df | Pr(Shapiro-Wilks) |
|---|---|---|---|---|---|---|---|---|
| (Intercept) | 1.18000 | (0.5933 - 1.766) | 3.288e-06 | 47 | 6 | 0.76 | 44 | 0.02 |
| Secretor | -1.81000 | (-0.9251 - -2.695) | 3.976e-13 | | | | | |
| log(DPP) | 0.01147 | (0.009961 - 0.01298) | 0.8642 | | | | | |

$GEE(z(log(\mathbf{DSLNT}+\epsilon)) \sim log(DPP) + Secretor$,id=subject,corstr='exchangeable')

| | Coef | 95% CI | Pr(Wald) | N. Obs | N. Groups | Marginal $R^2$ | df | Pr(Shapiro-Wilks) |
|---|---|---|---|---|---|---|---|---|
| (Intercept) | 0.7765 | (0.285 - 1.268) | 0.01619 | 47 | 6 | 0.34 | 44 | 0.22 |
| Secretor | 0.1627 | (0.1216 - 0.2038) | 0.2067 | | | | | |
| log(DPP) | -0.4691 | (-0.3124 - -0.6257) | 5.899e-3 | | | | | |

$GEE(z(log(\mathbf{DSLNH}+\epsilon)) \sim log(DPP) + Secretor$,id=subject,corstr='exchangeable')

| | Coef | 95% CI | Pr(Wald) | N. Obs | N. Groups | Marginal $R^2$ | df | Pr(Shapiro-Wilks) |
|---|---|---|---|---|---|---|---|---|
| (Intercept) | -0.7406 | (-0.5004 - -0.9808) | 7.600e-06 | 47 | 6 | 0.36 | 44 | 0.07 |
| Secretor | -0.2254 | (-0.1115 - -0.3393) | 0.3820 | | | | | |
| log(DPP) | 0.4740 | (0.3884 - 0.5597) | 2.682e-07 | | | | | |

**b.** Gaussian GEE, predicting motif abundance from DPP split on secretor status

$GEE(z(log(\mathbf{X62} + \epsilon)) \sim log(DPP) + Secretor$,id=subject,corstr='exchangeable',data=**'just-secretors')**

| | Coef | 95% CI | Pr(Wald) | N. Obs | N. Groups | Marginal $R^2$ | df | Pr(Shapiro-Wilks) |
|---|---|---|---|---|---|---|---|---|
| (Intercept) | -0.742 | (-0.48819 - -0.9957) | 2.12e-05 | 31 | 4 | 0.268 | 29 | 0.605 |
| log(DPP) | 0.399 | (0.33284 - 0.46529) | 2.44e-06 | | | | | |

$GEE(z(log(\mathbf{X62} + \epsilon)) \sim log(DPP) + Secretor$,id=subject,corstr='exchangeable',data=**'just-non-secretors'**)

| | Coef | 95% CI | Pr(Wald) | N. Obs | N. Groups | Marginal $R^2$ | df | Pr(Shapiro-Wilks) |
|---|---|---|---|---|---|---|---|---|
| (Intercept) | 1.218 | (1.0705 - 1.3654) | $< 2e-16$ | 16 | 2 | 0.687 | 14 | 0.881 |
| log(DPP) | -0.657 | (-0.63058 - -0.6832) | $< 2e-16$ | | | | | |

$GEE(z(log(\mathbf{LSTb} + \epsilon)) \sim log(DPP) + Secretor$,id=subject,corstr='exchangeable',data=**'just-secretors')**

| | Coef | 95% CI | Pr(Wald) | N. Obs | N. Groups | Marginal $R^2$ | df | Pr(Shapiro-Wilks) |
|---|---|---|---|---|---|---|---|---|
| (Intercept) | -0.393 | (-0.13891 - -0.64786) | 0.233 | 31 | 4 | 0.0884 | 29 | 0.9928 |
| log(DPP) | 0.217 | (0.17095 - 0.26372) | 0.046 | | | | | |

$GEE(z(log(\mathbf{LSTb} + \epsilon)) \sim log(DPP) + Secretor$,id=subject,corstr='exchangeable',data=**'just-non-secretors'**)

| | Coef | 95% CI | Pr(Wald) | N. Obs | N. Groups | Marginal $R^2$ | df | Pr(Shapiro-Wilks) |
|---|---|---|---|---|---|---|---|---|
| (Intercept) | 0.666 | (-0.11446 - 1.4473) | 0.264959 | 16 | 2 | 0.206 | 14 | 0.928 |
| log(DPP) | -0.359 | (-0.28515 - -0.43372) | 0.000653 | | | | | |

$GEE(z(log(\mathbf{DSLNT} + \epsilon)) \sim log(DPP) + Secretor$,id=subject,corstr='exchangeable',data=**'just-secretors')**

| | Coef | 95% CI | Pr(Wald) | N. Obs | N. Groups | Marginal $R^2$ | df | Pr(Shapiro-Wilks) |
|---|---|---|---|---|---|---|---|---|
| (Intercept) | 0.743 | (0.16746 - 1.3195) | 0.060 | 31 | 4 | 0.2237 | 29 | 0.0205 |
| log(DPP) | -0.389 | (-0.17451 - -0.60313) | 0.167 | | | | | |

$GEE(z(log(\mathbf{DSLNT} + \epsilon)) \sim log(DPP) + Secretor$,id=subject,corstr='exchangeable',data=**'just-non-secretors'**)

| | Coef | 95% CI | Pr(Wald) | N. Obs | N. Groups | Marginal $R^2$ | df | Pr(Shapiro-Wilks) |
|---|---|---|---|---|---|---|---|---|
| (Intercept) | 1.081 | (0.90883 - 1.2538) | $< 2e-16$ | 16 | 2 | 0.541 | 14 | 0.222 |
| log(DPP) | -0.583 | (-0.55665 - -0.60976) | $< 2e-16$ | | | | | |

$GEE(z(log(\mathbf{DSLNH} + \epsilon)) \sim log(DPP) + Secretor$,id=subject,corstr='exchangeable',data=**'just-secretors')**

| | Coef | 95% CI | Pr(Wald) | N. Obs | N. Groups | Marginal $R^2$ | df | Pr(Shapiro-Wilks) |
|---|---|---|---|---|---|---|---|---|
| (Intercept) | -0.993 | (-0.42991 - -1.5561) | 5.99e-04 | 31 | 4 | 0.448 | 29 | 0.245 |
| log(DPP) | 0.528 | (0.39235 - 0.66399) | 5.68e-05 | | | | | |

$GEE(z(log(\mathbf{DSLNH} + \epsilon)) \sim log(DPP) + Secretor$,id=subject,corstr='exchangeable',data=**'just-non-secretors'**)

| | Coef | 95% CI | Pr(Wald) | N. Obs | N. Groups | Marginal $R^2$ | df | Pr(Shapiro-Wilks) |
|---|---|---|---|---|---|---|---|---|
| (Intercept) | -0.662 | (-0.55101 - -0.77302) | 1.01e-14 | 16 | 2 | 0.2028 | 14 | 0.0203 |
| log(DPP) | 0.357 | (0.312 - 0.4021) | 2.91e-08 | | | | | |

**c.** Logistic GEE, predicting secretor status from estimated flux while controlling for DPP

$GEE(logit(Secretor) \sim log(DPP) + \mathbf{X62/X40}$ ,id=subject,corstr='exchangeable')

| | Coef | 95% CI | Pr(Wald) | N. Obs | N. Groups | Marginal Entropy | df |
|---|---|---|---|---|---|---|---|
| (Intercept) | 2.001310 | (-1.39521 - 5.39784) | 0.4230 | 47 | 6 | 0.50 | 44 |
| log(DPP) | 0.999898 | (0.999763 - 1.00003) | 0.1417 | | | | |
| I(X62/X40) | 0.989874 | (0.957091 - 1.02266) | 0.5469 | | | | |

$GEE(logit(Secretor) \sim log(DPP) + \mathbf{X106/X62}$ ,id=subject,corstr='exchangeable')

| | Coef | 95% CI | Pr(Wald) | N. Obs | N. Groups | Marginal Entropy | df |
|---|---|---|---|---|---|---|---|
| (Intercept) | 2.071930 | (-1.48709 - 5.63095) | 0.4058 | 47 | 6 | 0.50 | 44 |
| log(DPP) | 0.999989 | (0.998646 - 1.00133) | 0.9873 | | | | |
| I(X106/X62) | 0.948740 | (0.907258 - 0.990221) | 0.0183 | | | | |

**Supplementary Figure 1 | The glycoprofile clustering table with the original glycans.**

This is the clustering of sixteen glycoprofiles based on glycans. Since most of the glycans only exist in a few glycoprofiles, so the clustering mainly focus on the present/absent of the glycans (clusters 3-10), which means the information of structural similarity tend to be ignored in the clustering.

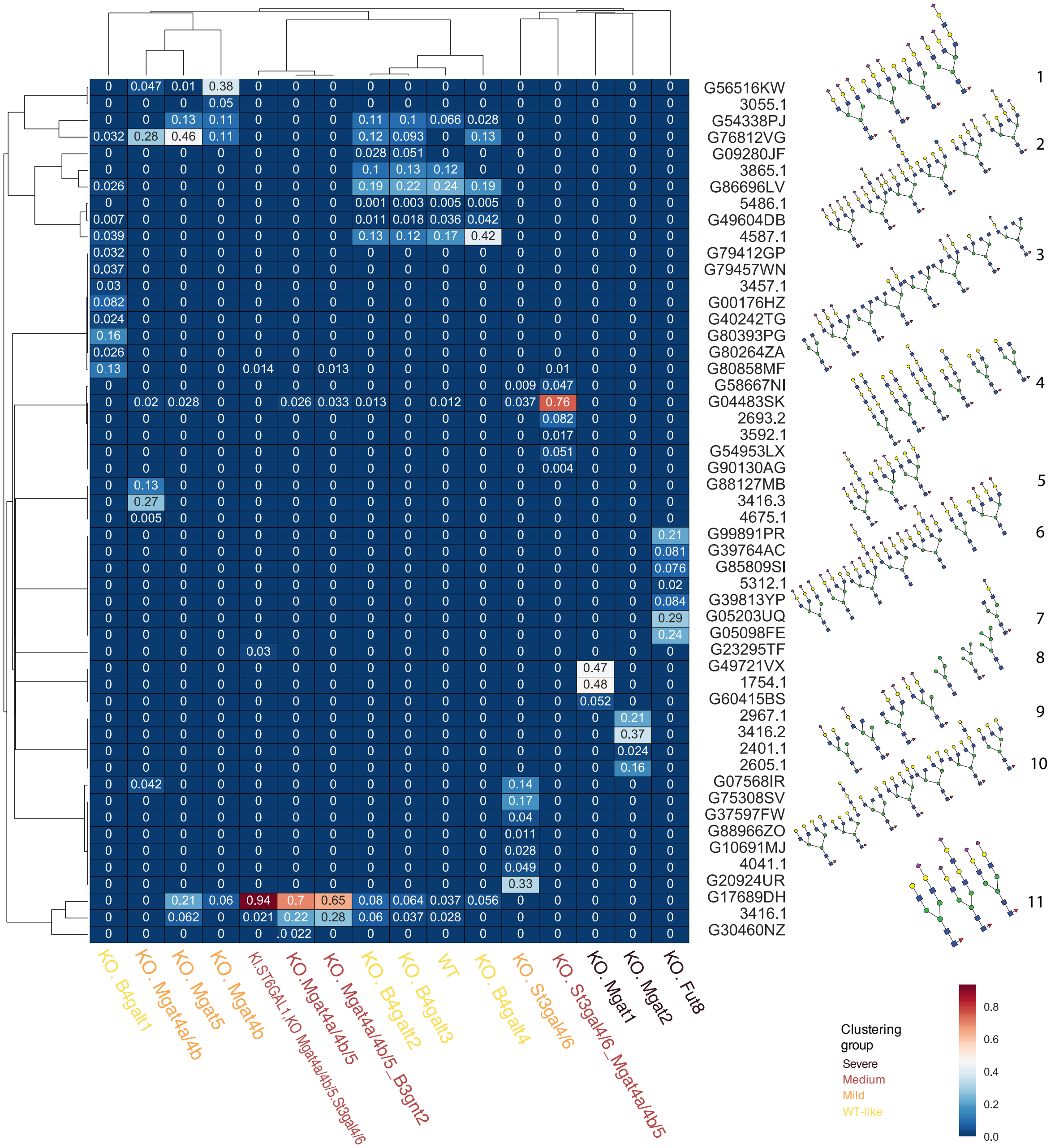

**Supplementary Figure 2 | Sample N-glycosylation network adapted from Spahn et al. 2016**

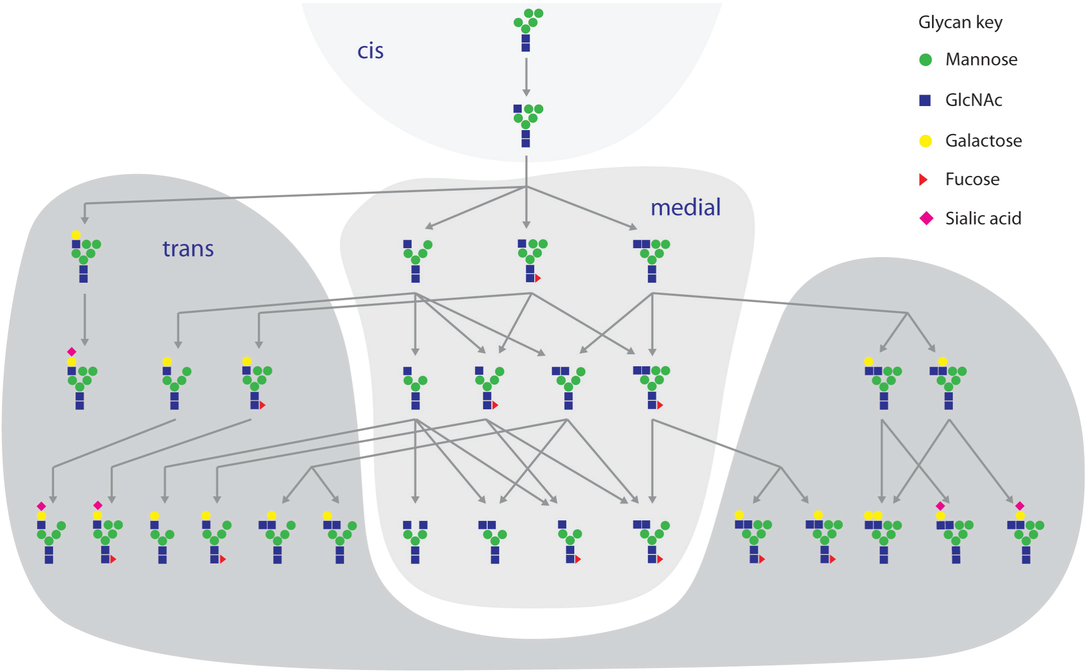


This is a theoretical N-glycan synthesized network. Our substructure network, starting with root, is able to mimic this network and is shown in **Supplementary Fig. 6**.

Spahn, P. N., Hansen, A. H., Hansen, H. G., Arnsdorf, J., Kildegaard, H. F., & Lewis, N. E. (2016). A Markov chain model for N-linked protein glycosylation–towards a low-parameter tool for model-driven glycoengineering. *Metabolic engineering*, *33*, 52-66.

Supplementary **Figure 3 | HMO substructure network with dependent substructure removed**

All the glycol-motifs are shown, and redundant nodes are merged. This is a directed-acyclic-graph and the direction goes from top to bottom. An edge with black color is important edge after merging that indicates the abundance changes. An edge with blue color is an edge that exists before merging that indicates the abundance variation between two substructures. The red lines are edges with 100% correlation but might not exists in the synthetic pathway because of the non-existent synthetic rules.

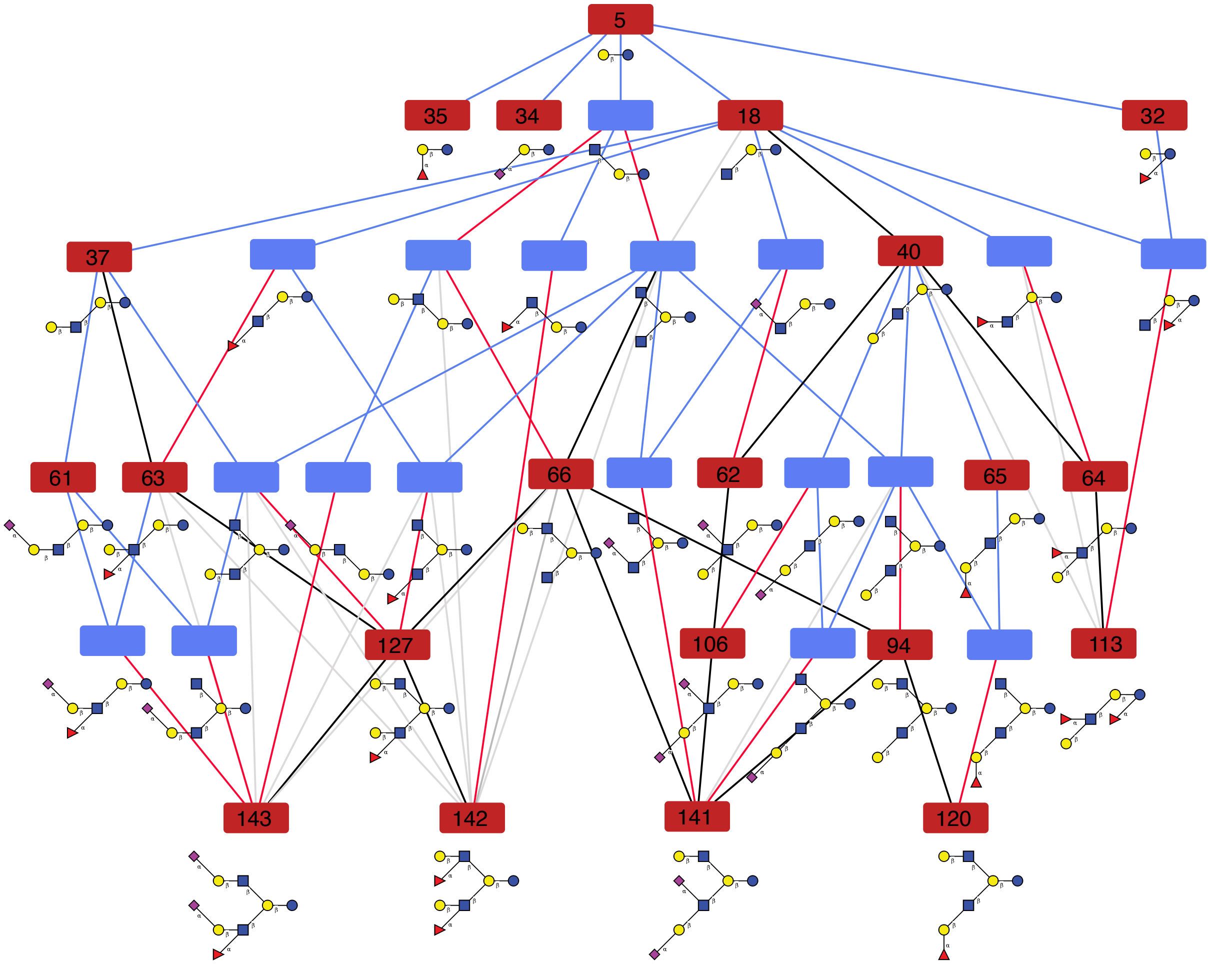

### Supplementary Figure 4 | Robustness of glyco-motifs clusters

This is the cluster of glyco-motif vector for EPO data. The robustness gives the criteria of much many substructure clusters should be generated. The cluster that has p-values > 0.95 are selected based on the p-value and the big block is further breakdown. We get 24 clusters in our EPO data.

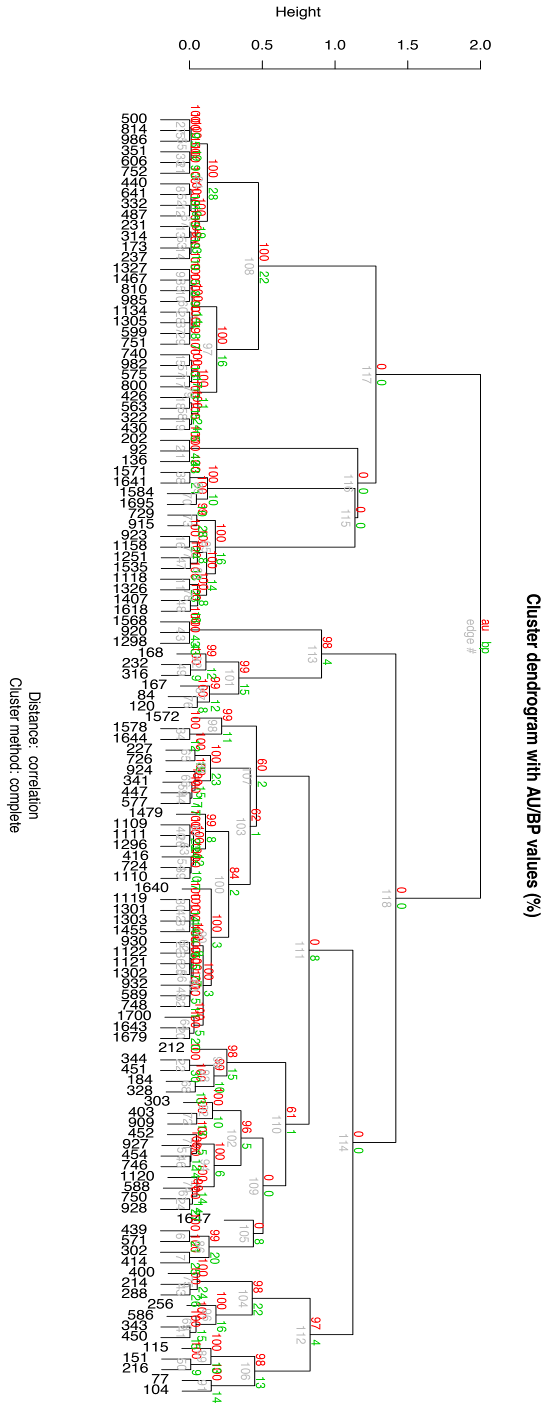

**Supplementary Figure 5 | The clustering robustness.**

The robustness is measured with BP (Bootstrap Probability) value. The BP values demonstrated the robustness. The clustering with glyco-motifs shows higher robustness than clustering with glycans. In the glycan clusters, the wild-like glycoprofiles are closer to the joint-knockouts that produces only biantennaries rather than the single Mgat4/5 knockouts that produces triantennaries which conflicts with the biological sense.

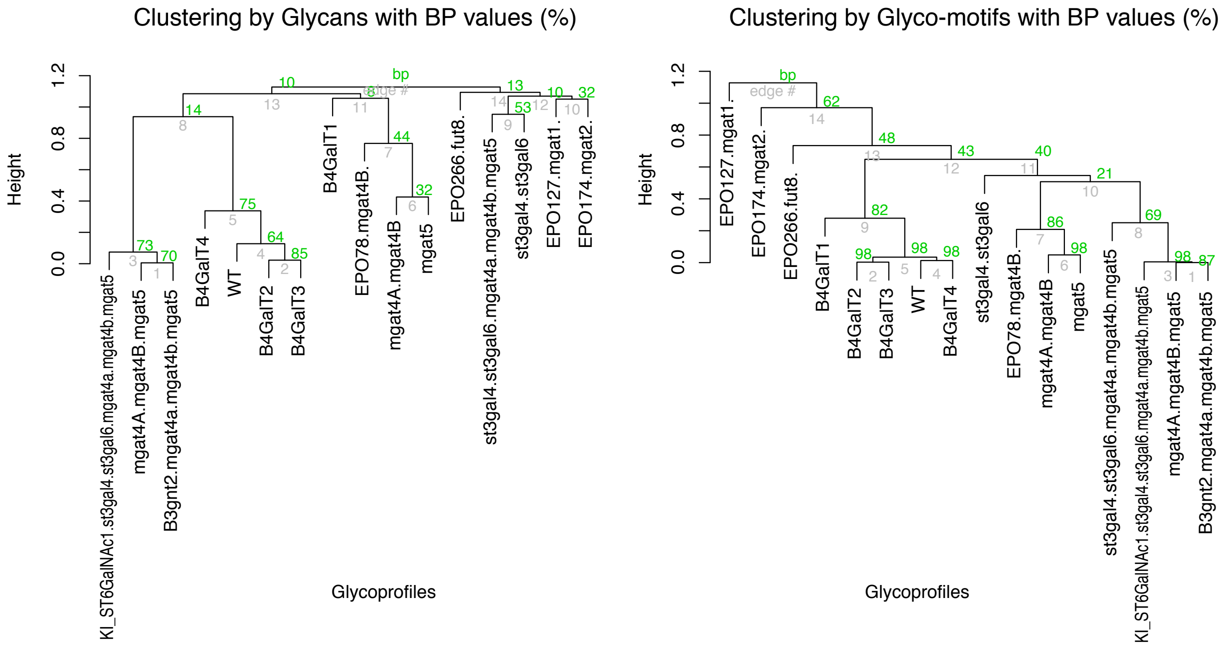

**Supplementary Figure 6 | The substructure network of EPO dataset**

The merged substructure network from 16 glycoprofiles contains 613 glyco-substructures. The red nodes are 120 glyco-motifs preserved. The light grey nodes are nodes can be filtered out. A dark grey node has many red child nodes which decrease its importance, so it can be removed. The red line indicates the abundances between child and parents node changed significantly so both nodes should be preserved. The blue lines indicate the abundances don't change significantly and the node above can be collapsed.

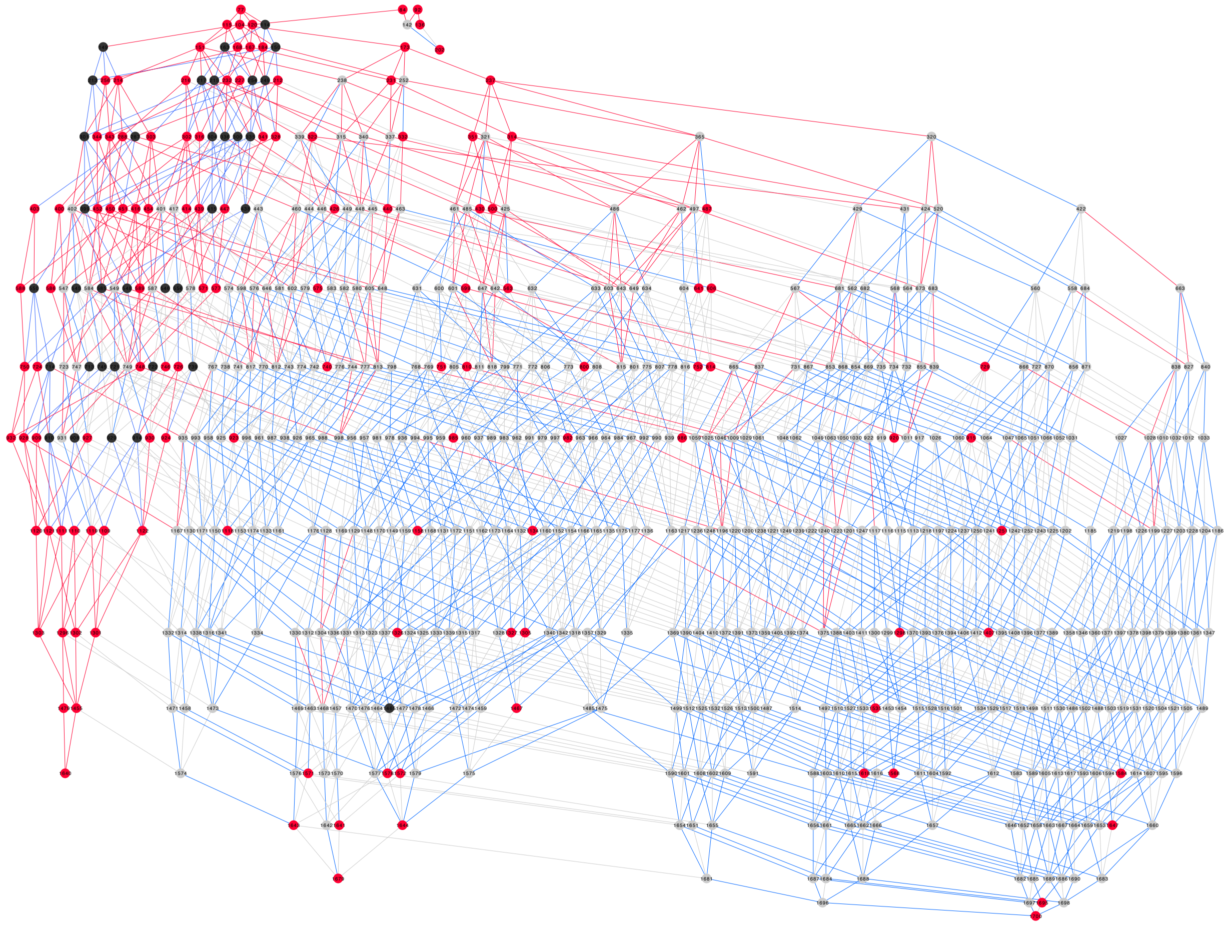

**Supplementary Figure 7 | Profile matching between the data from Čaval et al. (2018) and GlyCompare**

The lightened names are the knockouts that don't have MALDI-TOF data published. While some glyco-motif clusters can be seen in the glycoprofile clusters, there are important differences, and the glyco-motif clusters provide more information and improved cluster stability. Furthermore, the clustering result based on the glyco-motif was consistent with the clustering based on the native mass spectrometry, except for the Mgat2 knockout and the Fut8 knockout, which considerably changed the glycoprofiles by removing many common glycans. The main reason is that GlyCompare accounts for structural differences caused by each glycosyltransferase. This allows us to evaluate the magnitude of differences between glycans, whether it be between glycans with the same mass but different structural topologies, or subtle structural variations due to single changes in monosaccharides. Therefore, we had a better interpretation of the glycan structure variants across multiple glycoprofiles. All these results demonstrated the excellent performance of our GlyCompare in assessing the structural similarity between different glycoprofiles.

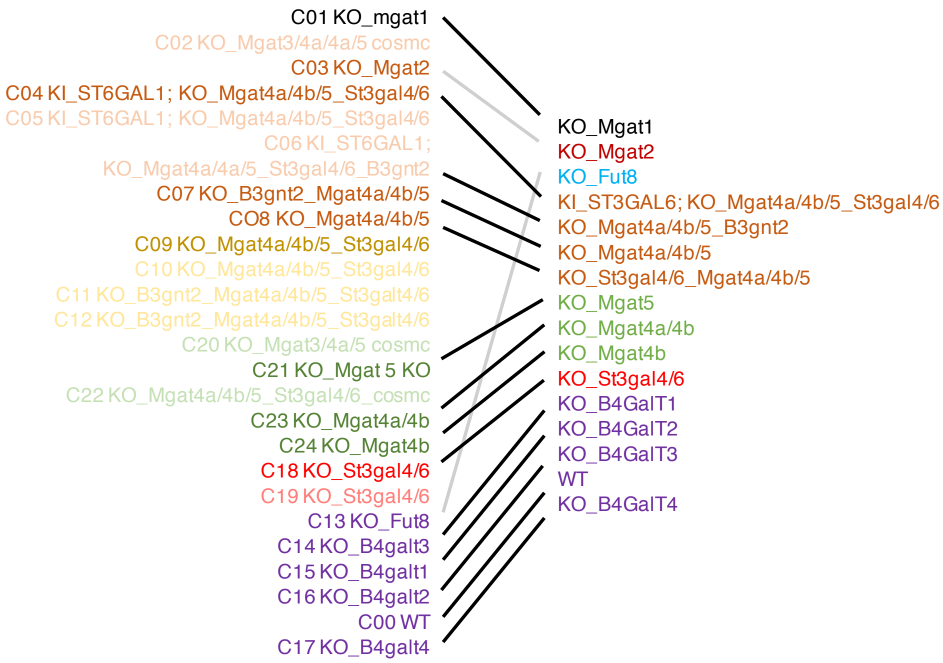


Čaval, T., Tian, W., Yang, Z., Clausen, H. & Heck, A. J. R. Direct quality control of glycoengineered erythropoietin variants. *Nat. Commun.* **9**, 3342 (2018).

**Supplementary Figure 8 | HMO dataset, the clustering of HMO by glycan using pearson correlation distance**

At the glycan-level, 2-fucosyllactose (2' FL) is the most abundant HMO in secretor mothers while Lacto-N-tetraose (LNT) and LNFPI are the most abundant HMOs in non-secretor mothers. The second major source of variance, DPP, shows a decrease in non-secretor LNFPI. At the substructure level, the clustering recapitulated the results from the raw HMO profiles and the α-1,2 fucosylated substructures were significantly associated with secretor status. The 2'FL substructure (X35) and the LNFPI substructure (X65) are significantly more abundant in secretor milk (Wald $p=2.35 \times 10^{-25}$, Wald $p=5.1 \times 10^{-12}$ respectively). The substructure abundance successfully reproduces the strongest effects known to be associated with secretor status.

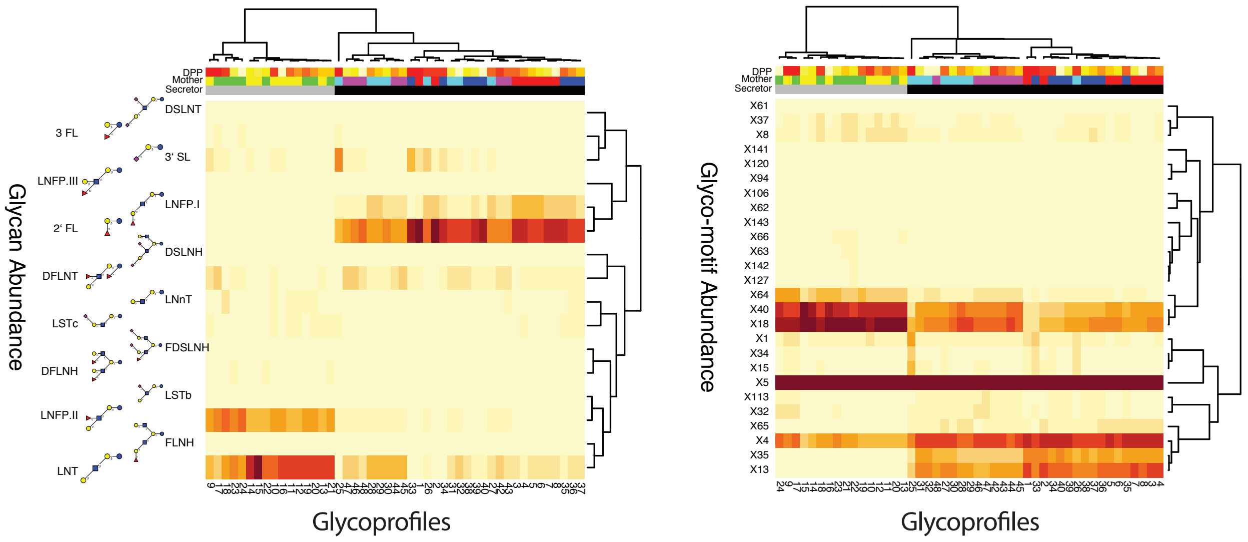

**Supplementary Figure 9 | HMO substructure general estimate equation coefficient and p-value plot**

Summary of regressions predicting either glycan or motif abundance from Days Postpartum (DPP) and Secretor status. The horizontal axis indicates the coefficient associating either DPP or secretor status with abundance and the vertical axis indicates the significance of that coefficient using the Wald test. Regression models were fit using Generalized Estimating Equations (GEE) with an exchangeable covariance structure to control for dependency structures within mothers. Colors indicate the identification of the glycan or glycan motif, size indicates the significance of the coefficient and shape indicate if the coefficient was attributed to DPP or secretor status. Models fit to predict glycan abundance (left) were of the form: GEE(z(log(S+ $\epsilon$))) ~ DPP + secretor, while models to predict motif abundance were of the form: GEE(z(log(S+ $\epsilon$))) ~ DPP + secretor. Where, z(x) is a z-score normalization to center and standardize abundance and $\epsilon = 0.001$. This link function was chosen because it fit a normal distribution (**Supplemental Table 3a)** and allowed for comparisons between regressions. There are some notable consistencies between the motif and glycan level results. As expected, 2'FL and its motif, X35, are both strongly and significantly enriched in secretor status. As are LNFPI and its motif, X65, are also strongly and significantly enriched with secretor status. Conversely, LSTb and X62 are negatively associated with secretor status. DPP has some significant but small associations negative associations with LSTc, 3'SL, and DSLNT. The 3'SL motif, X34 showed a consistent small negative significant association and the DSLNT motif. Most notably, X1, the sialic acid motif, was strongly negatively associated with DPP suggesting sialylation decreased in these sample over time.

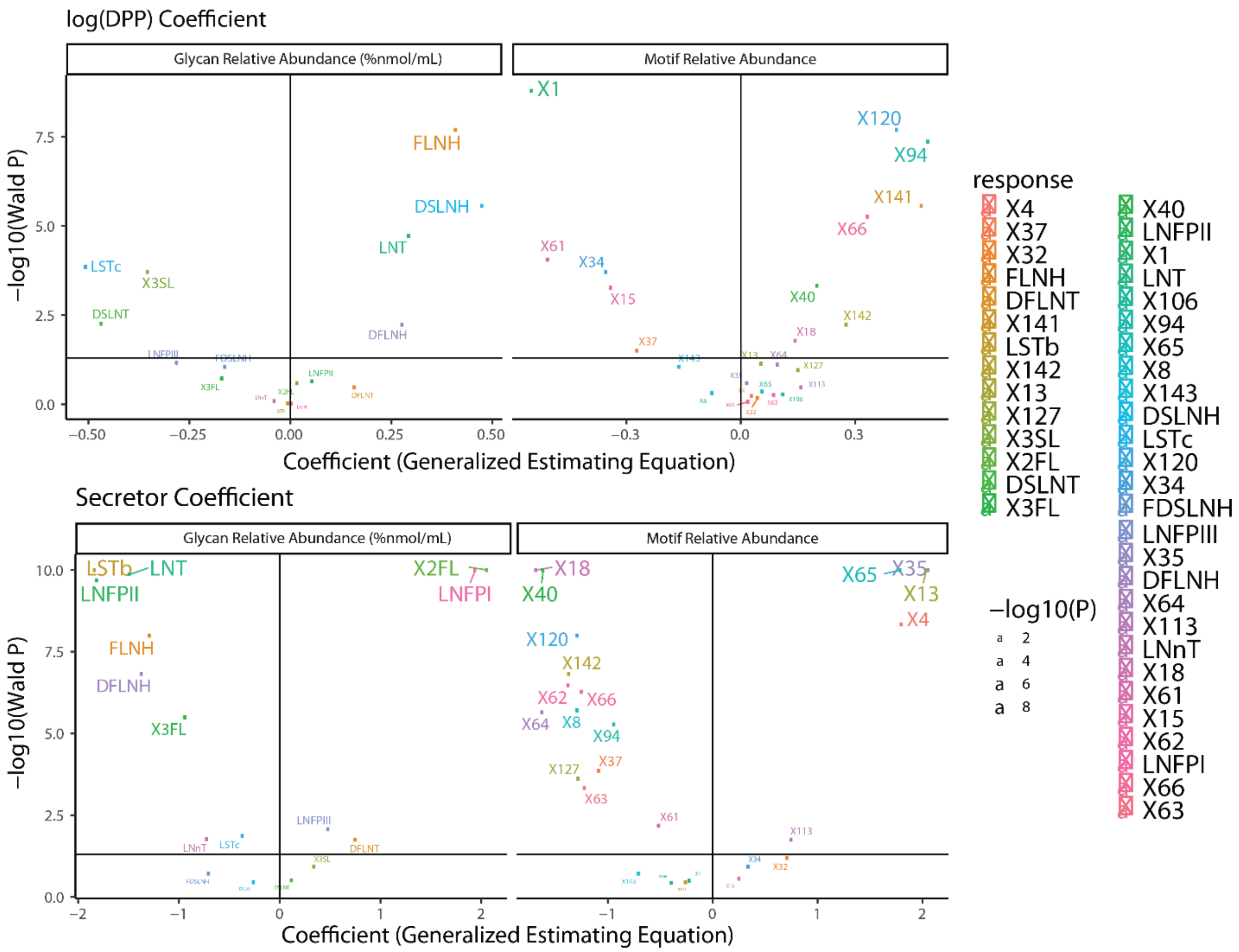
log(DPP) Coefficient
Glycan Relative Abundance (%nmol/mL)
Motif Relative Abundance
−log10(Wald P)
Coefficient (Generalized Estimating Equation)
FLNH
DSLNH
LNT
LSTc
X3SL
DSLNT
DFLNH
X1
X120
X94
X141
X66
X61
X34
X15
X40
X142
X18
X37
Secretor Coefficient
Glycan Relative Abundance (%nmol/mL)
Motif Relative Abundance
−log10(Wald P)
Coefficient (Generalized Estimating Equation)
LSTb
LNT
LNFPII
X2FL
LNFPI
FLNH
DFLNH
X3FL
LNFPIII
LNnT
LSTc
DFLNT
X18
X40
X65
X35
X13
X4
X120
X142
X62
X66
X64
X8
X94
X127
X37
X63
X61
X113
response
X4
X37
X32
FLNH
DFLNT
X141
LSTb
X142
X13
X127
X3SL
X2FL
DSLNT
X3FL
X40
LNFPII
X1
LNT
X106
X94
X65
X8
X143
DSLNH
LSTc
X120
X34
FDSLNH
LNFPIII
X35
DFLNH
X64
X113
LNnT
X18
X61
X15
X62
LNFPI
X66
X63
−log10(P)
2
4
6
8
